# The self-sufficient *Aspergillus fumigatus* Δ9-fatty acid desaturase SdeA requires a fused cytochrome b5 domain and hinge-dependent self-association for virulence

**DOI:** 10.64898/2026.10.23.760519

**Authors:** Jonatas Erick Maimoni Campanella, Gilberto Yanes Filho, João Henrique Tadini Marilhano Fabri, Natalia Fernanda Bueno, Paula Abrod Barbugli, Anderson Ferreira da Cunha, Endrews Delbaje, Leonardo Talachia Rosa, Iran Malavazi

**Author notes:** Centro de Biologia Molecular e Engenharia Genética, Universidade Estadual de Campinas (UNICAMP), Campinas, SP, Brazil. Corresponding author: Iran Malavazi.

## Abstract

Fungal Δ9-fatty acid desaturases are self-sufficient enzymes in which the electron-donating cytochrome b5 (Cytb5) module is fused to the catalytic core, yet whether this covalent linkage is architecturally required has remained untested. Here we show that in the human pathogen *Aspergillus fumigatus* the Cytb5 module of the Δ9-desaturase SdeA must act in *cis*, within a self-associated complex organized by a hinge segment. AlphaFold3 modeling predicts a dimer in which the Cytb5 domain of one protomer approaches the other’s di-iron center at a distance compatible with electron tunneling, and reciprocal co-immunoprecipitation shows that full-length SdeA self-associates *in vivo*. The recombinant Cytb5 module displays the spectral signatures of a genuine heme-binding redox domain. Nevertheless, deleting the Cytb5 domain, substituting its heme-coordinating histidines, or removing the 25-residue hinge each abolishes growth and virulence in *Galleria mellonella* without disrupting endoplasmic reticulum targeting, and a detached Cytb5-containing segment does not restore function. Hinge deletion abolishes stable SdeA self-association, identifying this segment as a structural organizer of the catalytically competent complex. Finally, we find no evidence that the *sdeB* locus provides a backup Δ9-desaturase. These results define the architectural requirements of a self-sufficient fungal desaturase and identify vulnerabilities absent from the human Δ9-desaturase for antifungal targeting.

## Introduction

Invasive fungal infections remain a major global health problem and are associated with substantial mortality, particularly in immunocompromised patients ^1, 2, 3^. Among the most clinically important filamentous fungal pathogens, *Aspergillus fumigatus* is the leading cause of invasive pulmonary aspergillosis and is also responsible for a wider range of chronic and allergic diseases ^4, 5, 6^. Its success as a pathogen relies on an exceptional capacity to adapt to host-associated stresses, including hypoxia, nutrient limitation, metal deprivation, immune attack, and antifungal exposure ^5^. As a result, aspergillosis remains difficult to manage clinically, reflecting the combined challenges of delayed diagnosis, complex host-pathogen interactions, and a limited antifungal arsenal^3, 7^.

Current antifungal therapy relies largely on compounds targeting the fungal cell wall or membrane, yet treatment options remain constrained by toxicity, limited success in some clinical settings, and the increasing emergence of resistance ^7, 8, 9, 10^. These limitations have reinforced the need to identify new membrane-associated vulnerabilities beyond ergosterol biosynthesis. In this context, lipid homeostasis has proven to be a vital determinant of fungal viability and pathogenicity. In particular, the biosynthesis of unsaturated fatty acids (UFA) by Δ9-fatty acid desaturases is necessary for maintaining membrane composition, fluidity, and function ^11, 12, 13, 14, 15^. By converting palmitic and stearic acids into palmitoleic and oleic acids, respectively, these enzymes sustain the *de novo* monounsaturated fatty acid (MUFA) pool required for fungal membrane integrity, helping maintain the balance between unsaturated and saturated fatty acids that underpins membrane fluidity, permeability, and adaptation to environmental stress. This pathway, therefore, represents an attractive yet still incompletely explored target for antifungal intervention, either as a standalone vulnerability or as a source of synergy with existing therapies.

In *A. fumigatus*, this essential desaturation step is catalyzed by SdeA, an endoplasmic reticulum (ER)-resident Δ9-fatty acid desaturase ^13, 15^. Genetic perturbation of *sdeA* profoundly disrupts fungal physiology, leading to auxotrophy for exogenous UFAs, altered membrane lipid composition, defective adaptation to temperature stress, and reduced virulence, underscoring the central role of this enzyme in fungal fitness and host adaptation ^13, 15, 16^. Mechanistically, the Δ9-fatty acid desaturation process relies on oxygen-dependent redox chemistry involving activation at a non-heme di-iron center at the catalytic module and sequential electron and proton transfer. This results in a cis double bond and water as products ^12, 17, 18, 19, 20^. Because Δ9-desaturation depends on the delivery of reducing equivalents to the di-iron catalytic center, SdeA must be organized to enable productive electron transfer to the active site. These considerations identify SdeA as both a critical metabolic node and a candidate membrane-associated vulnerability, while also exposing a central unresolved question: how this essential fungal desaturase is structurally organized to support the redox chemistry required for effective electron transfer and catalysis.

In contrast to mammalian SCD1, which relies on interactions with separate cytochrome b5 (Cytb5) and cytochrome b5 reductase (Cytb5R) proteins to complete electron transfer, fungal Δ9-fatty acid desaturases feature a unique self-sufficient fusion architecture with the Cytb5 domain positioned at the C-terminus of the same polypeptide chain ^12, 20^. Notably, *Saccharomyces cerevisiae* Ole1 was characterized with a fused Cytb5-like domain three decades ago ^21^. However, the structural and mechanistic implications of this architecture have remained largely unresolved, particularly in pathogenic fungi. This distinctive organization implies a fundamentally different mode of catalytic assembly, yet how the fused Cytb5 module is spatially arranged relative to the desaturase core to enable effective electron transfer to the catalytic center, culminating in the formation of the cis double bond, remains unknown. More fundamentally, whether the covalent linkage itself is required has remained untested: fusion could simply guarantee co-expression of two redox partners at fixed stoichiometry, or it could impose a geometry that a free-standing Cytb5 cannot reproduce.

To examine these unresolved structural and mechanistic questions, here we combine structural modeling, targeted mutagenesis, microscopic analyses, biochemical and invertebrate infection assays to dissect the molecular organization of the *A. fumigatus* Δ9-fatty acid desaturase SdeA. We show that SdeA is a Cytb5-dependent dimeric fusion enzyme in which the fused redox module cannot be functionally replaced by a detached copy, identify the heme-coordinating histidines essential for activity, and uncover a hinge region that enables self-association of the full-length SdeA and is essential for Cytb5-dependent function. We further address the unresolved question of whether *A. fumigatus* expresses a second functional Δ9-fatty acid desaturase by examining the annotation, transcriptional evidence, and functional behavior of the putative *sdeB* locus. Our study defines the architectural requirements of SdeA-dependent Δ9-fatty acid desaturation in a major human fungal pathogen and highlights fungal-specific features of this enzyme, including the fused Cytb5 module and hinge region, as candidate vulnerabilities for antifungal target prioritization.

## Results

### Structural refinement and functional validation identify SdeA as a Cytb5-dependent dimeric fusion enzyme required for virulence

Our recent perspective proposed that SdeA is a fungal Δ9-fatty acid desaturase in which the catalytic desaturase module is naturally fused to a C-terminal Cytb5 domain ^12^. Here we refined the structural organization of this enzyme to directly guide experimental interrogation of its associated mechanisms. Previous modeling had already suggested that fungal SdeA differs fundamentally from mammalian SCD1 (Stearoyl-CoA desaturase) by integrating the electron-donor Cytb5 module within the same polypeptide chain and had raised the possibility that productive electron transfer may depend on intermolecular rather than purely intramolecular domain organization ^12^. To address this question, we generated an AlphaFold3-based dimer model of full-length *A. fumigatus* SdeA and examined the relative positioning of its catalytic (fatty acid desaturation domain) and redox (Cytb5 domain) modules depicted in Fig. 1A. The resulting structure supports a defined dimeric arrangement in which the N-terminal desaturase core of one protomer (residues 19-297) is positioned in close apposition to the C-terminal Cytb5 domain of the adjacent protomer (Fig. 1B). The dimer model was obtained with pTM 0.75 and ipTM 0.72, indicating a well-defined fold and moderate confidence in the predicted interface. This spatial organization was the most plausible conformation that allows the interaction between the N-terminal desaturase and Cytb5 domain instead of a self-folding monomer ^12^. Structural alignment of SdeA with mammalian SCD1 (PDB 4ZYO) positioned stearoyl-CoA within the predicted catalytic pocket, while the heme prosthetic group of the neighboring Cytb5 domain localized near the desaturase active site in a configuration compatible with electron transfer in trans (Fig. 1C). In this arrangement, the heme iron of the Cytb5 domain lies 10.8-11.1 Å from the di-iron center of the opposing protomer, and the di-iron center lies 4.3-5.5 Å from the C9-C10 bond of the aligned substrate (Fig. 1C). The former separation falls within the ∼14 Å window over which physiological electron tunnelling operates in biological redox systems ^22^; measured edge-to-edge, the separation is shorter still. The predicted trans geometry is therefore chemically compatible with productive electron transfer. The model also preserved the canonical structural features expected for a membrane Δ9-desaturase catalytic center. Nine conserved histidines clustered within the desaturase core in positions corresponding to the metal-coordinating residues described for mammalian SCD1, supporting conservation of the di-metal-dependent desaturation chemistry in the fungal enzyme (Fig. 1B-C; yellow ribbons). At the same time, the Cytb5-containing C-terminal region (Leu 331-Arg409) displayed a well-defined heme-binding fold in which H366 and H392 are positioned as the axial ligands of the heme protoporphyrin iron (Fig. 1D, cyan). These features are relevant because they identify discrete structural elements within the fused Cytb5 module that are predicted to be essential for electron transfer, thereby providing direct targets for functional validation. To obtain independent biochemical support for this prediction, we expressed and purified the recombinant Cytb5 region of SdeA (Asp323-Ala456) (Fig. 1E) and analyzed its spectral properties using UV-visible spectroscopy ^23,24^. In the oxidized state (Fe^3+^), the purified protein displayed a prominent Soret absorption peak centered at 413 nm, a characteristic spectral feature of heme-containing proteins arising from the porphyrin ring and consistent with purification of the recombinant Cytb5 module as a heme-loaded holoprotein. Upon reduction with sodium dithionite, the Soret band shifted to 424 nm, accompanied by the appearance of two minor peaks at 527 nm and 557 nm, which are characteristic of the reduced form (Fe^2+^) of heme-bound cytochrome b5-like proteins (Fig. 1F). Thus, the structural model is supported not only by the predicted heme-binding geometry of H366 and H392, but also by biochemical evidence that the SdeA C-terminal module behaves as a genuine heme-binding, redox-active Cytb5 domain.

**Figure 1.**
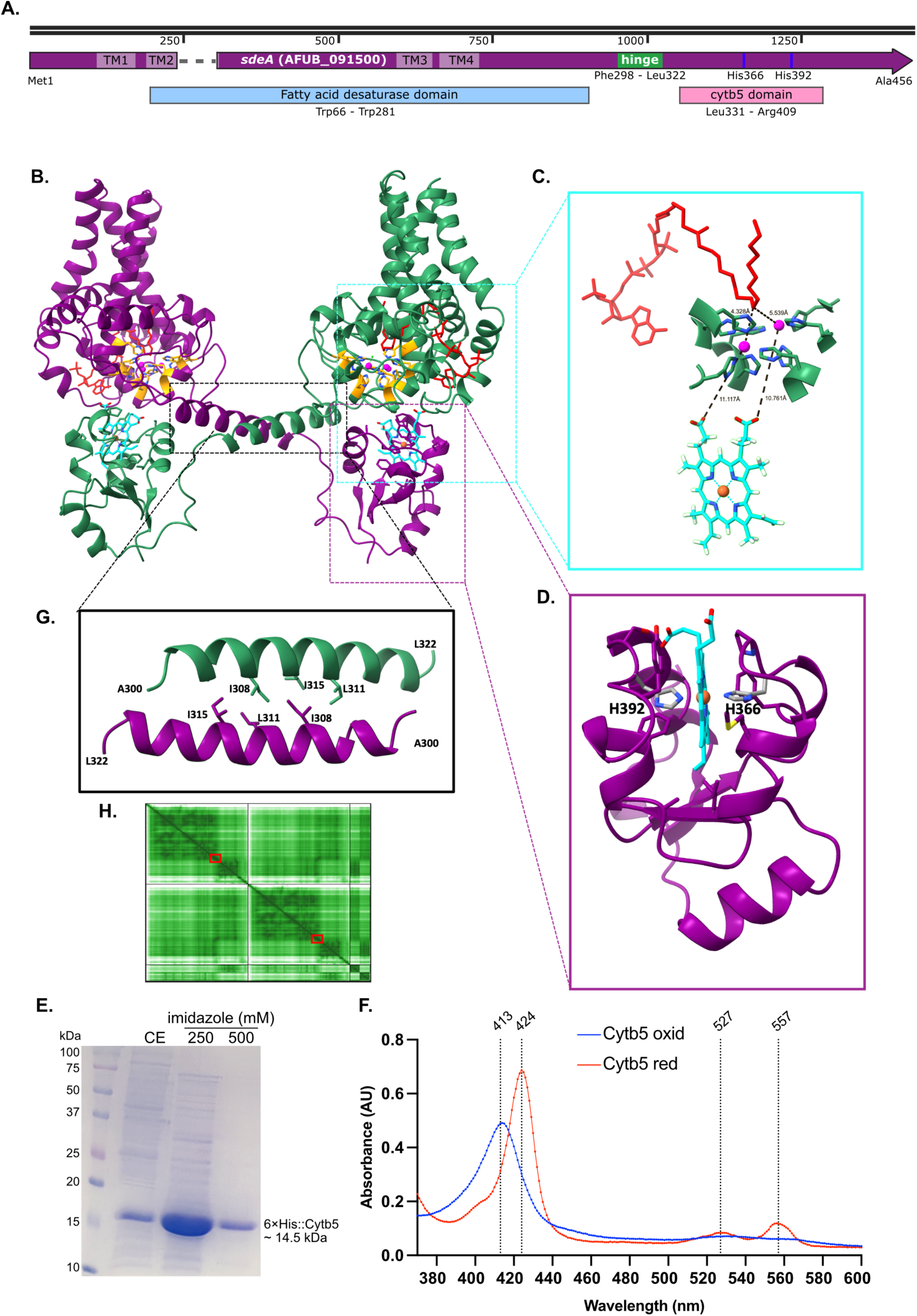
Structural model of SdeA supports a dimeric organization and a putative *trans* electron-transfer configuration. (A) Linear domain organization of *A. fumigatus* SdeA. The protein contains four predicted transmembrane segments (TM1-TM4), a fatty acid desaturase domain, an extended hinge region, and a C-terminal Cytb5 domain. The positions of the conserved heme-coordinating histidines H366 and H392 within the Cytb5 domain are indicated. (B) AlphaFold3-predicted model of the SdeA dimer (pTM 0.75; ipTM 0.72). The two SdeA protomers are shown in purple and green. The nine histidines predicted to coordinate Fe^3+^ ions within the desaturase core are shown in gold, with Fe^3+^ ions in magenta. The heme prosthetic group of the cytochrome b5 (Cytb5) domain is shown in cyan, with the central iron atom in orange. A stearoyl-CoA ligand, positioned by structural alignment of SdeA with human SCD1 (PDB: 4ZYO), is shown in red. The model supports a defined orientation between protomers, linked by an extended α-helical hinge region, and places the Cytb5 domain of one subunit in proximity to the catalytic region of the opposing protomer, consistent with a putative *trans* electron-transfer arrangement. (C) Enlarged view of the predicted interface between the Cytb5 heme group and the catalytic desaturase domain. The spatial relationship between the heme cofactor and the substrate-containing catalytic pocket supports a structurally coupled mechanism for electron delivery during fatty acid desaturation. Dashed lines show the distance between the heme group and Fe^3+^ ions (10.8-11.1 Å), as well as the ions and the double bond to be desaturated (4.3-5.5 Å). (D) Close-up of the Cytb5 domain (residues 331-409) showing the heme group coordinated by the conserved histidines H366 and H392, which are predicted to be required for iron coordination and electron transfer. Carbon and nitrogen atoms are depicted in gray and blue, respectively. (E) Expression and purification of the recombinant 6×His::Cytb5 fragment used for spectroscopic analyses. Representative SDS-PAGE showing the crude extract (CE) and immobilized metal affinity chromatography (IMAC) fractions. (F) UV-visible absorption spectra of purified recombinant Cytb5 in the oxidized and sodium dithionite-reduced states. The oxidized protein displayed a Soret peak at 413 nm, whereas reduction shifted the Soret band to 424 nm and generated visible bands at 527 and 557 nm, consistent with a heme-bound cytochrome b5-like redox domain. Data in E-F are representative of three independent experiments. (G) Detailed view of the hinge region (Phe298-Leu322), which connects the two major SdeA modules. The hydrophobic residues positioned along the opposing helices (residues A300-L322) are shown, highlighting a putative interaction surface that may stabilize protomer orientation. (H) Predicted aligned error (PAE) plot of the SdeA dimer model. Red boxes indicate the hinge regions. The overall confidence pattern supports stable intramolecular folding and a defined intermolecular arrangement within the dimer model.

A second unanticipated feature emerging from this refined model was the presence of an extended 25-amino-acid helical hinge (residues Phe298 to Leu322) connecting the desaturase and Cytb5 modules. In contrast to the earlier monomer-centered view, the dimer model interacts in an antiparallel fashion and places this segment at the interface between the two protomers (Fig. 1B, depicted in purple and green), where a series of hydrophobic side chains is positioned in a manner consistent with stabilizing protomer orientation (Fig. 1G). The predicted aligned error map further supported a defined intermolecular arrangement in the dimer model, including the hinge-containing region (Fig. 1H; red squares). These observations suggested that the hinge is unlikely to function merely as an inert linker. Instead, its position and predicted contacts raised the possibility that it actively contributes to the spatial register required to align the catalytic desaturase region with the redox-active Cytb5 domain within the prototypical fungal Δ9-desaturase enzyme family. This predicted refined model suggests a specific higher order of organization in which fusion and dimerization may act together to generate a catalytically productive architecture.

To test the first prediction arising from the structural model, we asked whether the fused Cytb5 module is required for SdeA function *in vivo*. In particular, we sought to address a fundamental mechanistic question: can the detached Cytb5-containing C-terminal segment or another Cytb5-containing protein functionally complement the SdeA desaturase core when the fused Cytb5 module is removed? For this purpose, we generated the *sdeA*^Δcytb^^5^::GFP allele, in which the C-terminal region containing the Cytb5 domain (Fig. 1A; Leu331 to Ala456) was removed while preserving the remainder of the protein (Met1-Gln330), and inserted this construct into the *aft4* Safe haven (Sh) locus ^25^ of the conditional *xylP::sdeA* background (Fig. S1). The Sh locus was chosen because it has been validated for ectopic gene expression in *A. fumigatus* and because a full-length *sdeA* allele placed there under its own promoter and terminator yields a stable, ER-targeted protein that self-associates with endogenously expressed SdeA (see below). We adopted this strategy after repeated unsuccessful attempts to generate *sdeA*^Δcytb5^ directly at the endogenous *sdeA* locus using a substitution cassette, a result consistent with the functional importance of this region. When endogenous *sdeA* expression was repressed by glucose, the *sdeA*^Δcytb5^::GFP strain was unable to grow, indicating that the truncated allele is unable to sustain SdeA activity under these conditions (Fig. 2A). Thus, the structural prediction that the Cytb5 domain is an integral and functionally unique determinant of SdeA function was strongly supported by genetic depletion of this module. We next asked whether the detached Cytb5-containing C-terminal segment could restore function in trans. To address this question, we expressed an ectopic version of the *cytb5* sequence under the control of the native *sdeA* promoter and fused to mRFP (Fig. S2) in the *xylP::sdeA* conditional mutant containing the *sdeA*^Δcytb5^::GFP at the Sh locus to generate an ectopic Cytb5+ strain. The resulting strain combines three experimental features: the endogenous *sdeA* locus remains under control of the repressible *xylP* promoter, the Sh carries the truncated sdeA^Δcytb5^::GFP allele, and an ectopic cassette expresses the detached Cytb5-containing segment fused to mRFP. Therefore, under repressing conditions for *xylP::sdeA*, this experimental design enables direct testing of whether the detached Cytb5-containing segment can restore function to the truncated SdeA protein *in trans*. In this strain, we retained an upstream sequence containing the QKKID region, which lies within the hinge-derived sequence (Fig. S2) and has previously been proposed as the second ER localization signal ^15^. This design was intended to minimize the possibility that a lack of complementation would simply reflect mislocalization of the detached redox module because it could not reach the ER. Despite this design, ectopic expression of the detached Cytb5-containing segment did not restore growth upon glucose repression of endogenous *sdeA* (Fig. 2A). The phenotype of non-viability displayed by all the mutant strains employed was rescued by supplementation with palmitoleic acid or oleic acid, consistent with impaired Δ9-desaturation and consequent failure to maintain MUFA biosynthesis (Fig. 2A). Together, these data indicate that the Cytb5 region cannot rescue SdeA activity as an independently expressed module, consistent with a requirement for structural integration of the Cytb5 region within SdeA (Fig. 1).

**Figure 2.**
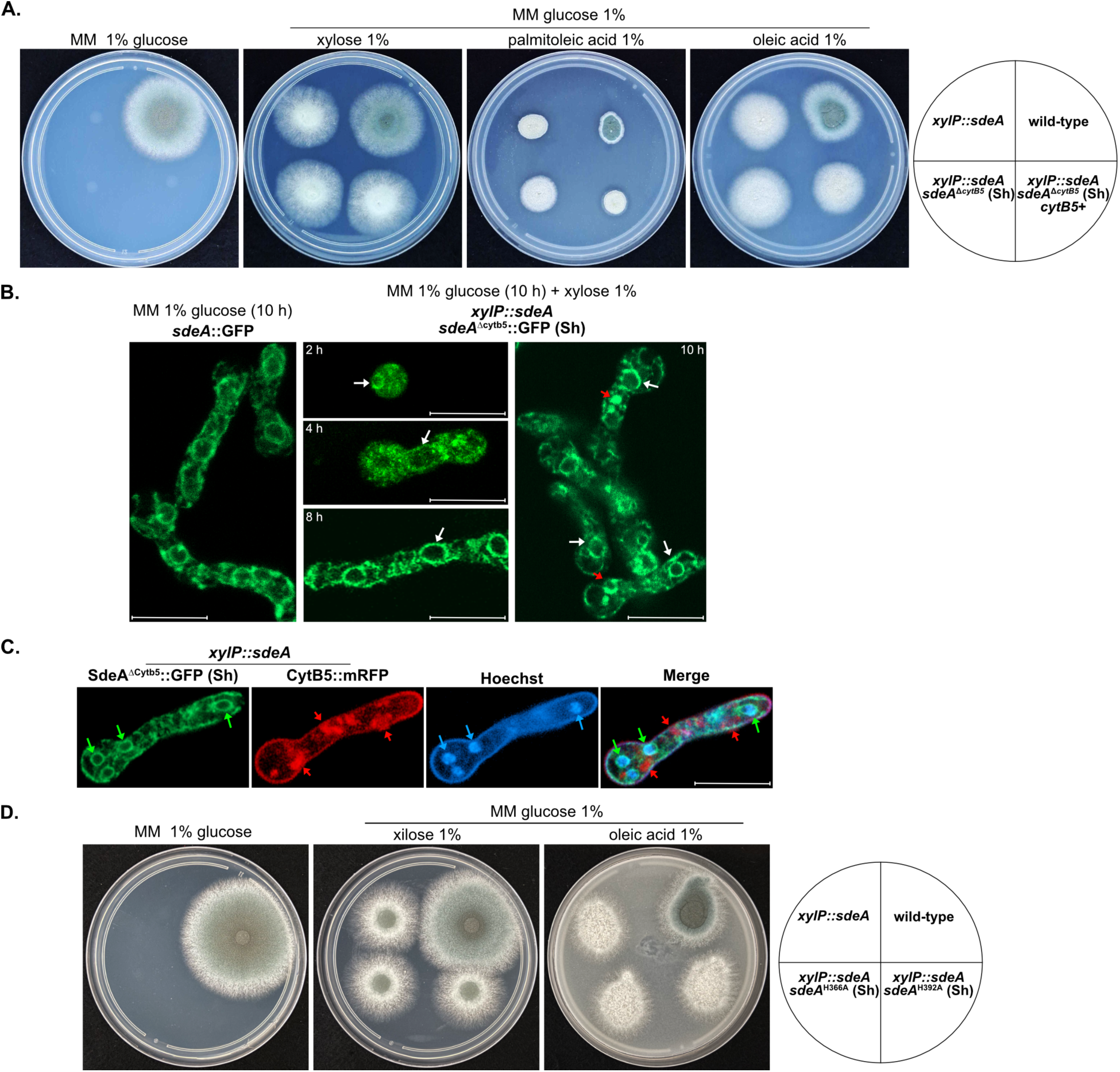
The Cytb5 domain and its conserved heme-coordinating histidines are essential for SdeA function and activity. (A) Growth assay testing the functional requirement of the Cytb5 domain and the ability of a detached Cytb5-containing segment to restore SdeA activity in *trans*. 1×10^4^ conidia of the wild-type strain, the parental *xylP::sdeA* mutant, the *xylP::sdeA; sdeA*^Δcytb5^::GFP (Sh) strain, and the *xylP::sdeA; sdeA*^Δcytb5^::GFP (Sh); Cytb5^+^ strain ectopically expressing Cytb5::mRFP were grown in MM under the indicated conditions for 60 h at 37°C. Glucose repression of *xylP::sdeA* revealed that the *sdeA*^Δcytb5^::GFP allele is non-functional under these conditions, and this defect was not complemented by ectopic expression of the detached C-terminal Cytb5-containing segment. Supplementation with palmitoleic acid or oleic acid restored the phenotype, consistent with defective unsaturated fatty acid biosynthesis upon loss of Cytb5-dependent SdeA activity. (B) Confocal microscopy analysis of subcellular localization of SdeA lacking the Cytb5 domain. Wild-type SdeA::GFP was examined under glucose conditions, whereas the *sdeA*^Δcytb5^::GFP strain was first maintained under glucose repression and then shifted to xylose for the indicated times before imaging. The truncated SdeA^ΔCytb5^::GFP protein retained a reticular fluorescence pattern consistent with ER localization (white arrows), indicating that deletion of the Cytb5 domain does not abolish ER targeting (Scale Bar, 10 µm). (C) Confocal analysis of the strain co-expressing *sdeA*^Δcytb5^::GFP and the detached C-terminal Cytb5-containing segment fused to mRFP under permissive conditions. Cells were grown in MM + xylose 1% for 8 h to allow expression of the conditional allele and to evaluate whether the ectopically expressed Cytb5-containing segment colocalizes with the truncated SdeA^Δcytb5^ protein. SdeA^Δcytb5^::GFP retained perinuclear ER-associated fluorescence adjacent to Hoechst-stained nuclei (green and blue arrows), whereas Cytb5::mRFP was detected mainly in punctate cytosolic structures consistent with protein aggregates (red arrows). (D) Growth assay evaluating the functional importance of the conserved Cytb5 histidines H366 and H392. Full-length mutant alleles carrying the substitutions H366A or H392A, integrated at the Sh locus under the endogenous *sdeA* promoter and terminator, were tested in the conditional *xylP::sdeA* background under glucose, xylose, and oleic acid supplementation (72 h). Neither mutant allele restored growth under glucose repression of endogenous *sdeA*, whereas oleic acid rescued the phenotype, indicating that both histidines are required for SdeA function and supporting their predicted role in heme coordination within the Cytb5 domain. Data in A-D are representative of three independent experiments.

We then examined whether loss of function in the *sdeA*^Δcytb5^::GFP mutant could be explained by defective subcellular targeting. Confocal microscopy showed that the truncated SdeA^Δcytb5^::GFP protein retained a wild-type reticular fluorescence pattern consistent with ER localization ^15^ after xylose reinduction to break dormancy (2-10 h) and allow the expression of the truncated SdeA^Δcytb5^ isoform within the cell (Fig. 2B). Thus, removal of the Cytb5 domain alone does not abolish ER targeting, indicating that the severe functional defect of the truncation mutant cannot be attributed simply to mislocalization of the protein. We also performed confocal analysis of the strain co-expressing SdeA^Δcytb5^::GFP together with the detached Cytb5::mRFP module under permissive conditions to determine whether the two proteins occupied overlapping subcellular regions (Fig. 2C). This analysis was particularly relevant because the isolated SdeA Cytb5 domain retained heme-dependent redox responsiveness *in vitro*, as shown by Soret spectral shift (Fig. 1F), indicating that this module can behave as a functional redox domain when expressed independently. Under the conditions tested, the Cytb5::mRFP signal did not detectably overlap with SdeA^Δcytb5^::GFP, indicating that the detached Cytb5-containing segment is not stably recruited to the same perinuclear ER-associated regions (green and blue arrows) as the truncated SdeA protein. Instead, Cytb5::mRFP was detected mainly in punctate cytosolic structures consistent with protein aggregates (red arrows). This lack of colocalization provides an additional explanation for the failure of the detached Cytb5 module to complement the growth defect, suggesting that productive SdeA activity requires more than independent expression of a redox-competent Cytb5 segment. Because the detached module did not detectably reach the ER, we cannot formally separate a requirement for cis integration of the Cytb5 domain from a failure of the isolated module to be correctly targeted. Both interpretations converge, however, on the conclusion that the redox module cannot function as an autonomous unit under these conditions.

The structural model further predicted that the Cytb5 heme-binding fold should depend critically on the conserved histidines H366 and H392, which are positioned as the axial ligands of the heme iron (Fig. 1D). To test this prediction directly, we generated full-length *sdeA*^H366A^ and *sdeA*^H392A^ alleles under the control of the endogenous promoter and terminator regions (Fig. S3) and inserted them into the Sh locus of the conditional *xylP::sdeA* strain to evaluate the ability of these mutated proteins to individually rescue growth under conditions of genetic depletion of the endogenous *sdeA*. In both cases, glucose-mediated repression of endogenous *sdeA* revealed a severe growth defect that was not rescued by the mutant alleles, whereas oleic acid supplementation restored growth (Fig. 2D). These results show that both histidines are indispensable for SdeA activity and strongly support the interpretation that the fused Cytb5 domain functions as an electron transfer module rather than as a passive structural appendage.

Because SdeA-dependent production of MUFAs is tightly linked to fungal fitness, we next asked whether the same architectural features required for growth are also required for virulence. In the *G. mellonella* infection model, the strain carrying the *sdeA*^Δcytb5^::GFP allele in the conditional *xylP::sdeA* background was strongly attenuated, and ectopic expression of the detached Cytb5-containing segment also failed to restore pathogenicity (Fig. 3A). Likewise, strains carrying the full-length *sdeA*^H366A^ or *sdeA*^H392A^ alleles remained avirulent in the same background (Fig. 3B). Thus, both the integrity of the fused Cytb5 domain and the heme-coordinating residues that define its electron transfer center are required for vegetative growth and virulence.

**Figure 3.**
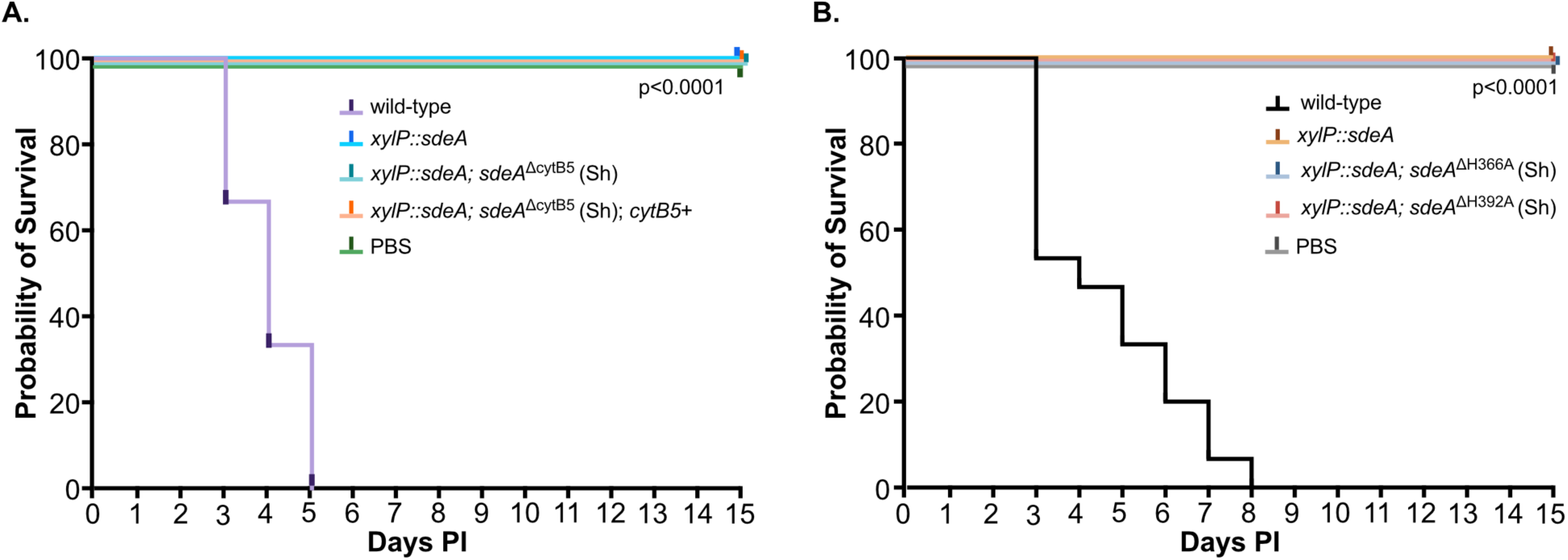
The Cytb5 domain and its heme-coordinating histidines are required for full virulence in *Galleria mellonella*. (A) Survival of *G. mellonella* larvae following infection with the indicated strains to assess the contribution of the Cytb5 domain to SdeA-dependent virulence. Fifteen larvae per group were infected with 1×10^6^ conidia per larva and monitored daily for 15 days post-infection (PI). Data are from three independent experiments. Deletion of the Cytb5 domain strongly attenuated virulence relative to the wild-type strain. Likewise, the strain carrying the *sdeA*^Δcytb5^ allele at the Sh locus in the conditional *xylP::sdeA* background remained severely attenuated, and ectopic expression of the detached Cytb5-containing segment fused to mRFP did not restore virulence. PBS-injected larvae were included as a control. (B) Survival of *G. mellonella* larvae infected with strains carrying full-length SdeA alleles mutated at the conserved Cytb5 histidines H366 or H392. Larvae were infected with 1×10^6^ conidia and followed for 15 days PI. Data are from three independent experiments. Both the *sdeA*^H366A^ and *sdeA*^H392A^ strains were strongly attenuated in virulence, indicating that these conserved residues are essential for SdeA function during infection. PBS-injected larvae served as the negative control. Survival curves were analyzed using the log-rank (Mantel-Cox) test in GraphPad Prism 10. P < 0.0001 for all mutants compared to the wild-type strain.

Taken together, the results in Figs. 2-3 provide functional validation of the predicted structural model proposed in Fig. 1. These findings indicate that the fused Cytb5 domain is required for SdeA function and that endogenous Cytb5-containing proteins present in the cell cannot compensate for its loss, since the sdeA^Δcytb5^ strain failed to grow despite their presence. A detached SdeA-derived Cytb5-containing segment likewise failed to restore function, although this segment did not detectably reach the ER. They also show that its conserved heme-coordinating histidines are necessary for SdeA-dependent growth and virulence. Collectively, these results position the fused Cytb5 module as a central functional determinant of the fungal enzyme and set the stage for testing whether the newly identified hinge region also contributes to assembly of the catalytically competent SdeA complex.

### The hinge region enables SdeA self-association and is essential for Cytb5-dependent function

Our structural prediction model identified a short hinge region linking the desaturase domain to the C-terminal Cytb5 domain as a previously unrecognized and potentially important architectural element of SdeA. This α-helix segment occupies a strategic position between the catalytic domain of the enzyme and the fused redox module and contains sequence features highlighted by the structural model, which suggest a possible contribution to intramolecular organization and interprotomer assembly (Fig. 1A-B and 1G). We therefore asked whether the hinge region is required for SdeA function *in vivo*. To this end, a hinge-deleted allele (Phe298 to Leu322) that preserved the *sdeA* reading frame was generated and inserted into the Sh locus in the conditional *xylP::sdeA* background (Fig. S4). Again, this strategy was employed after repeated unsuccessful attempts to generate *sdeA*^Δhinge^ directly at the endogenous *sdeA* locus using a substitution cassette, a result consistent with the functional importance of this region. In the presence of glucose that represses endogenous *sdeA*, the *sdeA*^Δhinge^ allele failed to support growth, demonstrating that this region is indispensable for SdeA activity (Fig. 4A). Supplementation with palmitoleic acid or oleic acid restored growth (Fig. 4A), indicating that the defect caused by hinge deletion is directly linked to impaired Δ9-desaturation and consequent failure to sustain MUFA biosynthesis. As observed for the deletion of the Cytb5 domain, removal of the hinge region renders SdeA non-functional in the absence of endogenous *sdeA* expression. To determine whether the loss of function caused by hinge deletion could be explained by defective subcellular targeting, we examined the localization of SdeA^Δhinge^::GFP by confocal microscopy. Wild-type SdeA::GFP displayed the expected reticular fluorescence pattern consistent with ER localization under glucose conditions (Fig. 4B). After initial glucose repression followed by xylose reinduction, SdeA^Δhinge^::GFP expressed from the Sh locus was also detected in reticular ER-associated structures, indicating that deletion of the hinge region does not abolish ER targeting. However, in contrast to wild-type SdeA::GFP, the hinge-deleted protein also accumulated in large punctate cytosolic structures consistent with protein aggregates (Fig. 4B). Thus, the severe growth defect of the sdeA^Δhinge^ allele cannot be attributed simply to loss of ER localization, but rather suggests that the hinge region is required for the proper structural organization and/or stability of SdeA within the ER membrane environment.

**Figure 4.**
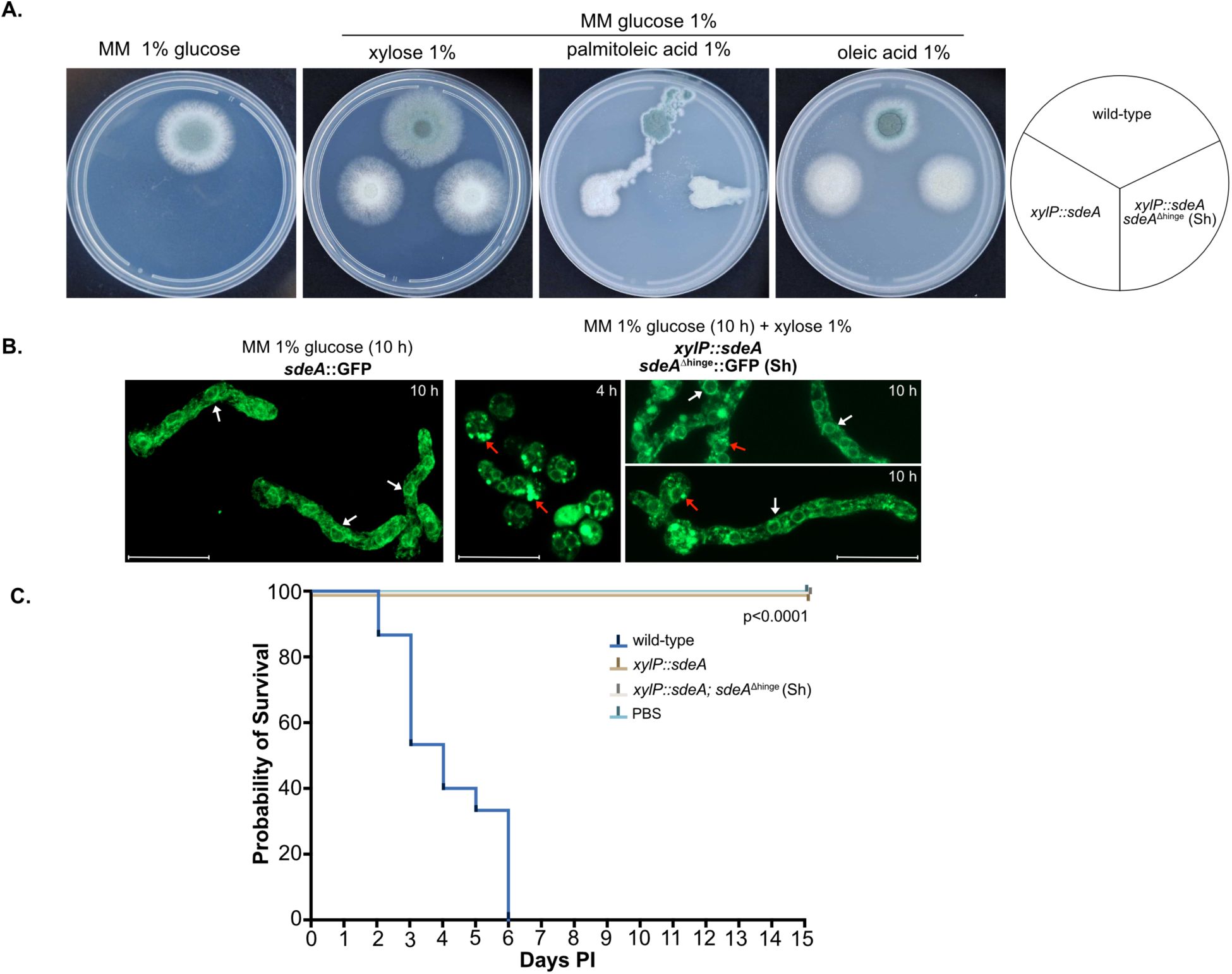
The hinge region is a key structural determinant of SdeA function and virulence. (A) Growth assay assessing the functional requirement of the hinge region linking the catalytic desaturase domain of SdeA to the C-terminal Cytb5 domain. 1×10^4^ conidia of the indicated strains were inoculated on MM supplemented with 1% glucose, 1% xylose, 1% palmitoleic acid, or 1% oleic acid and grown for 60 h at 37°C. When endogenous *sdeA* expression was repressed by glucose, the *sdeA*^Δhinge^ allele failed to support growth, demonstrating that the hinge region is required for SdeA activity. Growth was restored by supplementation with palmitoleic acid or oleic acid, consistent with impaired unsaturated fatty acid biosynthesis in the absence of an intact hinge region. (B) Confocal microscopy analysis of subcellular localization of SdeA lacking the hinge segment. Wild-type SdeA::GFP was examined under glucose conditions, whereas the *sdeA*^Δhinge^::GFP strain was first maintained under glucose repression for 10 h and then shifted to 1% xylose for the indicated times before imaging. The SdeA^Δhinge^::GFP protein expressed from the Sh retained a reticular fluorescence pattern consistent with ER localization (white arrows), although large punctated cytosolic structures consistent with protein aggregates were also detected (red arrows). Scale Bar, 10 µm. (C) Survival of *G. mellonella* larvae following infection with the indicated strains to assess the contribution of the hinge region to SdeA-dependent virulence. Fifteen larvae per group were infected with 1×10^6^ conidia and monitored daily for 15 days post-infection (PI). Data are from three independent experiments. Deletion of the 25-residue hinge region strongly attenuated virulence relative to the wild-type strain, indicating that this segment is required for SdeA function during infection. PBS-injected larvae were included as a control. Survival curves were analyzed using the log-rank (Mantel-Cox) test in GraphPad Prism 10. P < 0.0001 for all mutants compared to the wild-type strain. Data in A-B are representative of three independent experiments.

The requirement for the SdeA hinge region was also investigated in the context of the *G. mellonella* infection model. The data indicated that the *xylP::sdeA*; *sdeA*^Δhinge^ Sh strain was severely attenuated and failed to restore virulence relative to the wild-type strain (Fig. 4C), indicating that the hinge region is also essential for SdeA-dependent pathogenic fitness, suggesting that the 25-residue-long hinge is a key determinant of SdeA function during both vegetative growth and host infection. Because the hinge region occupies the junction between the catalytic desaturase module and the C-terminal Cytb5 domain in our structural model, we next investigated whether its essentiality might reflect a more fundamental role in organizing the active enzyme complex. In the AlphaFold3-based SdeA dimer model, the hinge contributes to the relative orientation of the two protomers and lies at the structural interface that positions the catalytic desaturase region of one monomer in register with the Cytb5-containing C-terminus of the other (Fig. 1B and 1G). This arrangement predicts that hinge-dependent dimerization is required to establish the geometry necessary for electron transfer from the fused Cytb5 module to the desaturase active site, thereby enabling conversion of saturated fatty acids into palmitoleic and oleic acids. We tested this prediction directly by asking whether SdeA self-associates *in vivo* and whether this interaction depends on the hinge region. In a strain co-expressing full-length SdeA::3×HA from the Sh locus and SdeA::GFP from the endogenous *sdeA* locus, reciprocal Co-IP recovered the partner-tagged protein in both directions (Fig. 5A-B), demonstrating that full-length SdeA undergoes self-association *in vivo*. These results provide experimental support for the dimeric organization predicted by the structural model. Notably, these experiments also show that a full-length *sdeA* allele placed at the Sh under its endogenous promoter and terminator yields a stable protein that is correctly targeted to the ER (Fig. 2B and 4B) and retains the capacity to self-associate with endogenous SdeA, arguing against the possibility that the loss of function observed for the Sh-expressed mutant alleles reflects the ectopic location of the constructs rather than the engineered lesions. The consequence of hinge deletion on this interaction was subsequently examined in a strain co-expressing full-length SdeA::3×HA together with SdeA^Δhinge^::GFP. In clear contrast to the full-length/full-length combination, reciprocal Co-IP failed to detect an association between full-length SdeA and the hinge-deleted variant (Fig. 5C-D), despite the efficient recovery of both baits in the corresponding direct-capture lanes. Thus, deletion of the hinge abolishes stable SdeA self-association.

**Figure 5.**
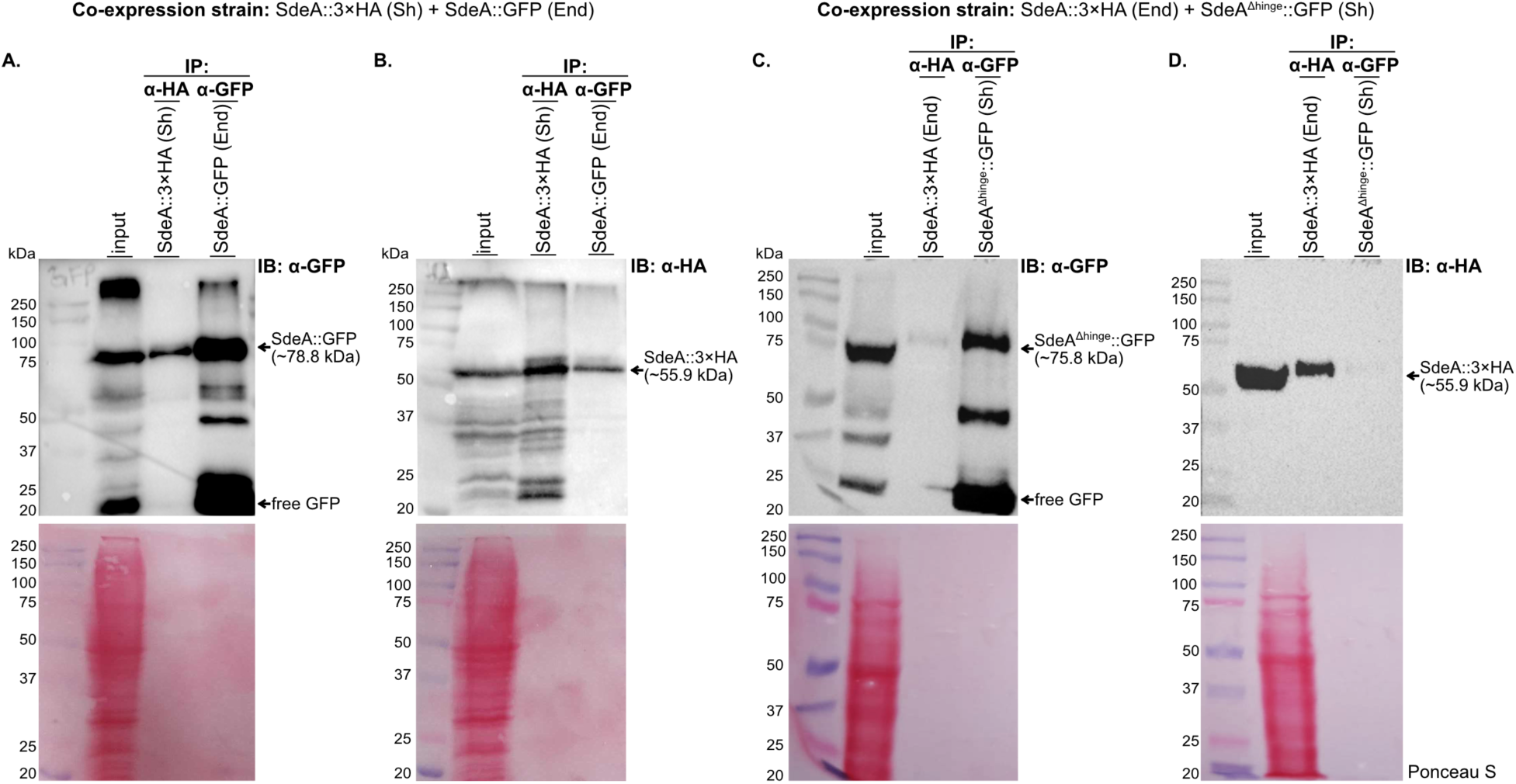
SdeA self-association depends on the hinge region. (A-B) Reciprocal co-immunoprecipitation assays showing self-association of full-length SdeA *in vivo*. A single strain co-expressing SdeA::3×HA from the Safe haven (Sh) locus and SdeA::GFP from the endogenous *sdeA* locus (End) was used for both assays. In every panel, the two reciprocal immunoprecipitates obtained from the same extract are resolved side by side and identified by the antibody used for capture (α-HA, bait SdeA::3×HA; α-GFP, bait SdeA::GFP); panels A and B therefore display the same pair of immunoprecipitations revealed with different antibodies. In each panel, the lane in which the immunoblotted tag matches the captured bait serves as a direct-capture control confirming that the corresponding resin recovered its target efficiently, whereas the other lane reports co-precipitation of the partner protein. In panel A, full-length SdeA::GFP was recovered in the α-HA immunoprecipitate, demonstrating co-precipitation with SdeA::3×HA. In panel B, the reciprocal experiment recovered SdeA::3×HA in the α-GFP immunoprecipitate. (C-D) Reciprocal co-immunoprecipitation assays testing the contribution of the hinge region to SdeA self-association. A single strain co-expressing SdeA::3×HA from the endogenous *sdeA* locus (End) and SdeA^Δhinge^::GFP from the Sh locus was used for both assays and processed exactly as in A-B. In C (IB: α-GFP), only a faint residual SdeA^Δhinge^::GFP signal was recovered in the α-HA immunoprecipitate, in sharp contrast to the robust co-precipitation of full-length SdeA::GFP observed in A. In D, SdeA::3×HA was not recovered in the α-GFP immunoprecipitate. In both panels, the direct-capture lanes confirm that each resin efficiently recovered its bait, establishing that the loss of co-precipitation reflects the absence of a stable interaction rather than a failed immunoprecipitation. Because C and D were performed with the same resins, extract amounts, and protocol as A and B, they additionally control for non-specific retention of SdeA on the affinity matrices. Together, these results indicate that deletion of the hinge region abolishes stable SdeA self-association. For all panels, input corresponds to total protein extract prior to immunoprecipitation. Sh: *aft4* safe-haven locus; End: endogenous *sdeA* locus. Ponceau S staining of the corresponding membranes is shown below as a transfer/control stain. The positions of the full-length GFP fusion proteins and the free GFP species are indicated in panels A and C. Immunoblots are representative of three independent experiments.

Taken together, our results support assigning a central mechanistic role to the hinge region, required not only for SdeA-dependent growth and virulence but also for the formation of the self-associated complex predicted by the three-dimensional model. These findings support a model in which the hinge is not an autonomous self-association motif but rather enables assembly of the catalytically competent SdeA complex, thereby positioning the desaturase catalytic region and the fused C-terminal Cytb5 module in the proper spatial arrangement for productive electron transfer and catalysis.

### A putative additional Δ9-desaturase locus (*sdeB*) is ambiguously annotated and fails to compensate for SdeA loss

Because *A. nidulans* encodes two functional Δ9-fatty acid desaturases, SdeA and SdeB ^26^, we asked whether *A. fumigatus* similarly harbors a second functional Δ9-desaturase capable of supporting MUFA biosynthesis. Additional searches for Δ9-desaturase-related loci in the *A. fumigatus* A1163 and Af293 genomes identified AFUB_090930 and Afu7g05350 as the most plausible *sdeB* candidates, respectively ^12^. We therefore sought to determine whether this locus could provide functional compensation for SdeA loss. A closer inspection of AFUB_090930 (*sdeB*^A1163^), which is annotated as a pseudogene, further indicated that the deposited sequence is disrupted by a thymine insertion at nucleotide position 200, which is predicted to shift the reading frame and introduce a premature stop codon, thereby generating a truncated putative SdeB protein of only 87 amino acids. However, Sanger sequencing of the corresponding *sdeB* genomic region in Af293, CEA10, CEA10 derivative strains (CEA17, Δ*akuB*^KU80^, and A1160), and the A1163 strain available in our laboratory showed that this insertion was absent from all strains analyzed (Fig. S5A). These results indicate that the T200 insertion is not a conserved feature of the *sdeB* locus across the strain backgrounds examined here, but also highlight inconsistencies among available genome annotations.

Because the putative *sdeB*^A1163^ locus showed inconsistencies among available annotations, we re-examined its genomic organization in the Af293 strain (*sdeB*^Af293^; not annotated as a pseudogene), using published RNA-seq datasets to assess whether the locus is transcriptionally represented in the *A. fumigatus* transcriptome and to infer potential intron-containing regions. Analysis of 13 publicly available RNA-seq datasets identified read coverage consistent with transcription across the *sdeB*^Af293^ locus, but also revealed discrepancies in the predicted intron structure. RNA-seq read alignments suggested a putative intron-containing region that does not match the intron annotation currently assigned to Afu7g05350 (Fig. S5B). However, based solely on these alignments and bioinformatics tools, we were unable to define the mature *sdeB* transcript sequence, resolve the precise exon-intron boundaries, or determine the putative polypeptide sequence. Notably, the putative intron region suggested by transcript alignment appears to be conserved in the close relative *A. fischeri* ortholog NFIA_026560, whereas the currently annotated *A. fumigatus* sequence appears to lack the region corresponding to the predicted C-terminal Cytb5-like domain present in *A. fischeri*, suggesting that this domain may be absent from the *A. fumigatus* SdeB protein (Fig. S5B), a possibility we examine at the protein level below. Thus, although the *sdeB* locus is not transcriptionally silent, its mature coding sequence remains unresolved.

Next, based on the available sequence and the predicted translation start site, which aligns with the corresponding start region of NFIA_026560, we investigated whether a putative SdeB^Af293^ protein product could be detected. To this end, we introduced an N-terminal V5 epitope into the *sdeB*^Af293^ allele and placed this construct under the control of the strong *gpdA* promoter at the Sh locus in the *xylP::sdeA* background. The N-terminal tag was chosen to minimize reliance on an uncertain predicted C-terminus for protein detection (Fig. S6A-B). Western blot analysis of this *gpdA*::V5::*sdeB*^Af293^ ectopic construct did not detect a V5-reactive signal corresponding to the molecular weight predicted for the intact *sdeB*^Af293^ product under the conditions tested. Instead, a low-molecular-weight V5-reactive species of approximately 20 kDa was observed (Fig. S5C). Given that the predicted full-length SdeB protein contains the conserved fatty acid desaturase domain and is expected to be substantially larger (∼40 kDa), the detected signal is unlikely to correspond to the intact protein. These results suggest that the expressed product may undergo extensive proteolytic processing, premature translation termination, or accumulation as a truncated polypeptide. Alternatively, and consistent with the RNA-seq evidence above (Fig. S5B), the discrepancy between the intron-exon structure inferred from read alignments and the current Afu7g05350 model raises the possibility that the mature SdeB protein is shorter than the ∼40 kDa predicted from the deposited annotation, in which case the ∼20 kDa species could reflect a genuinely truncated SdeB lacking the C-terminal region.

We next asked whether the *sdeB* locus could provide functional compensation for loss of *sdeA*. To this end, the *sdeB*^Af293^ allele was introduced into the Sh locus of the conditional *xylP::sdeA* strain under its putative endogenous 5’ and 3’ regulatory regions. Upon glucose-mediated repression of endogenous *sdeA*, this construct did not restore growth, and the same result was observed using the *gpdA*::V5::*sdeB*^Af293^ allele (Fig. 6A). As expected, growth was restored by supplementation with palmitoleic acid or oleic acid, reinforcing that the observed defect reflects loss of SdeA-dependent desaturation of palmitic and stearic acids, rather than a nonspecific growth defect. In the *G. mellonella* infection model, neither the Sh *sdeB*^Af293^ allele expressed from its endogenous regulatory elements nor the *gpdA*-driven *sdeB*^Af293^ construct restored virulence in the *xylP::sdeA* background (Fig. 6B). Thus, under the conditions tested here, the analyzed *sdeB* constructs cannot replace *sdeA* either for vegetative growth or pathogenic fitness. Because these constructs did not yield a detectable intact SdeB product, however, this result alone does not establish that a functional SdeB protein would be unable to compensate.

**Figure 6.**
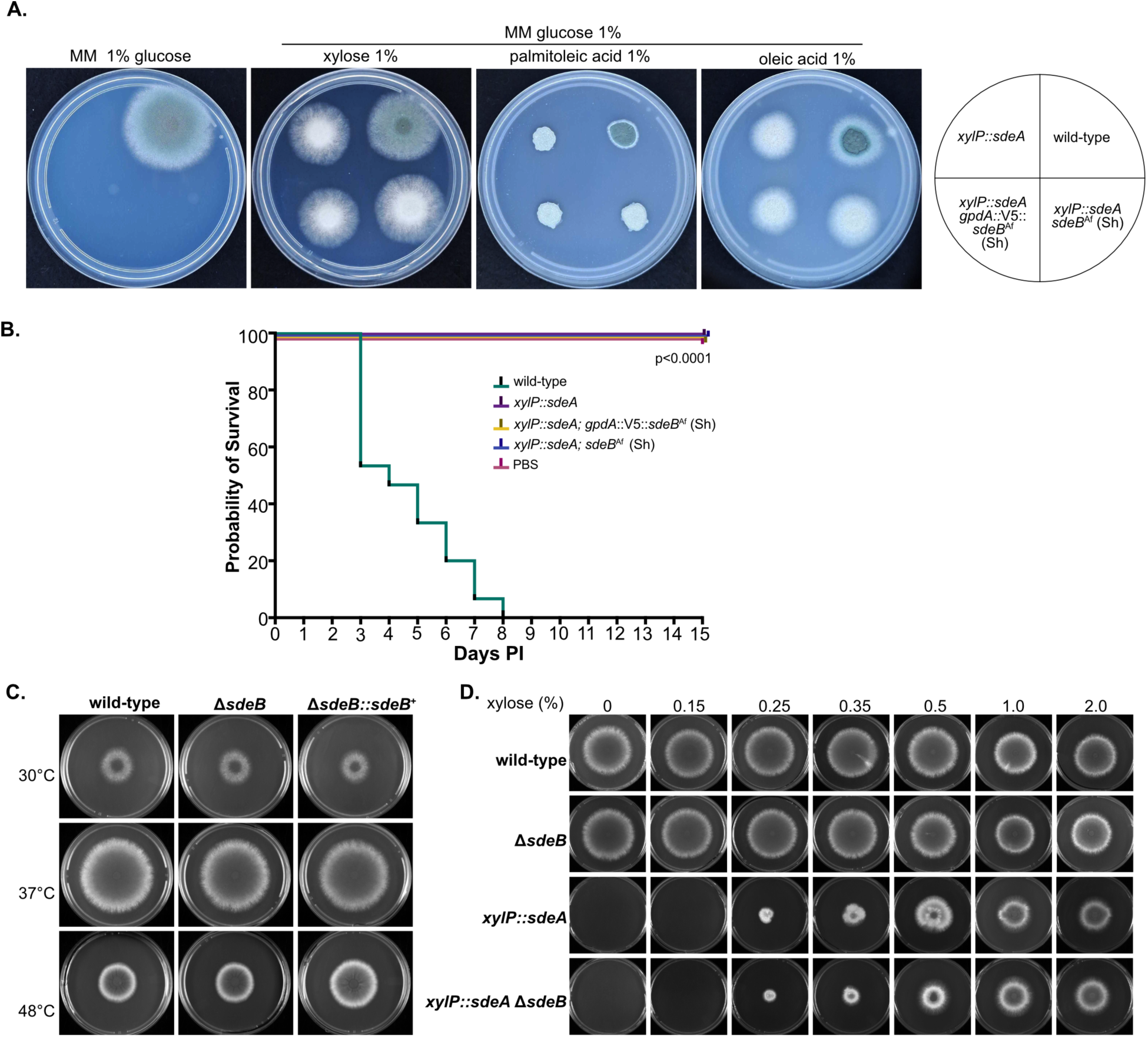
SdeB does not compensate for SdeA loss under the tested conditions. (A) Growth assay evaluating whether *sdeB* can functionally compensate for *sdeA* loss. The indicated strains, including *xylP::sdeA*; *sdeB* (Sh), *xylP::sdeA*; *gpdA*::V5::*sdeB* (Sh), the parental conditional mutant *xylP::sdeA,* and wild-type (1×10^4^ conidia), were grown on minimal medium supplemented with 1% glucose, 1% xylose, 1% palmitoleic acid, or 1% oleic acid. Repression of endogenous *sdeA* by glucose caused a marked growth defect that was not rescued by insertion of *sdeB* at the Sh locus under its endogenous promoter, nor by ectopic overexpression of *sdeB* from the *gpdA* promoter. Growth was restored by supplementation with palmitoleic acid or oleic acid, consistent with impaired MUFA biosynthesis upon loss of SdeA function. (B) Survival of *G. mellonella* larvae following infection with the indicated strains to assess whether SdeB restores SdeA-dependent virulence. Fifteen larvae per group were infected with 1×10^6^ conidia per larva and monitored daily for 15 days post-infection (PI). Data are from three independent experiments. Neither *sdeB* inserted at the Sh under its endogenous regulatory elements nor *gpdA*-driven *sdeB* overexpression restored virulence in the *xylP::sdeA* background. PBS-injected larvae were included as a control. Survival curves were analyzed using the log-rank (Mantel-Cox) test in GraphPad Prism 10. No significant difference was detected between the *sdeB*-expressing strains and wild-type or the parental *xylP::sdeA* strain (P > 0.05). (C) 1×10^4^ conidia of the wild-type, Δ*sdeB*, and Δ*sdeB*::*sdeB*^+^ strains were inoculated onto Petri dishes containing MM and incubated at 30, 37, or 48°C for 96 h. Under these conditions, deletion of *sdeB* did not cause an evident colony growth defect. (D) 1×10^4^ conidia of the wild-type, *xylP::sdeA*, *ΔsdeB*, and *xylP::sdeA* Δ*sdeB* strains were inoculated onto solid MM containing increasing concentrations of xylose, and plates were incubated at 37°C for 96 h. Deletion of *sdeB* did not aggravate the growth defect of the *xylP::sdeA* strain across the xylose gradient, indicating no detectable genetic interaction between *sdeA* and *sdeB* under the conditions tested. Data in A, C-D are representative of three independent experiments.

To test the contribution of the *sdeB* locus itself, independently of the tagged constructs, we deleted *sdeB* in the Δ*akuB*^KU80^ parental background and complemented the deletion (Fig. S6C-F). No obvious defects in colony growth were observed for the Δ*sdeB* mutant at 30, 37, or 48°C, and the complemented Δ*sdeB::sdeB*^+^ strain behaved as the parental strain (Fig. 6C). We then asked whether *sdeB* interacts genetically with *sdeA*. If *sdeB* encoded a Δ9-desaturase able to contribute to MUFA biosynthesis, its removal would be expected to exacerbate the growth defect caused by *sdeA* depletion. However, deletion of *sdeB* did not markedly aggravate the growth defect of the conditional *xylP::sdeA* strain across a xylose gradient (Fig. 6D). This argues against a backup Δ9-desaturase function for this locus under the conditions tested.

Together, these data indicate that *A. fumigatus* harbors an *sdeB*-related locus that is transcriptionally represented but ambiguously annotated, with unresolved exon-intron boundaries that preclude confident prediction of the mature polypeptide and its domain architecture. Most directly, deletion of the locus produced no growth phenotype and did not aggravate the defect caused by *sdeA* depletion, arguing that *sdeB* does not contribute detectably to MUFA biosynthesis under these conditions. The two *sdeB* expression constructs tested likewise restored neither growth nor virulence in the *xylP::sdeA* background, although the absence of a detectable intact SdeB product limits the interpretation of those experiments. These findings do not exclude the possibility that the *sdeB* locus may have a condition-specific role or produce an alternative product under circumstances not examined here. However, under the conditions in which SdeA-dependent Δ9-desaturation is required for MUFA biosynthesis, growth, and virulence, we find no evidence that *sdeB* functions as a backup for SdeA.

## Discussion

Fungal Δ9-fatty acid desaturases have emerged as central regulators of membrane homeostasis in pathogenic fungi ^11, 13, 14, 15^. In *A. fumigatus*, SdeA supports the production of the major MUFAs palmitoleic and oleic acids, yet the mechanistic basis by which this enzyme supports Δ9-desaturation remains poorly understood. Here, we show that SdeA is not simply a desaturase catalytic core bearing an accessory redox appendage, but an integrated fungal fusion enzyme in which the C-terminal Cytb5 domain, conserved heme-coordinating histidines, hinge-dependent self-association and dimeric organization cooperate to sustain Δ9-desaturase activity (Fig. 7).

**Figure 7.**
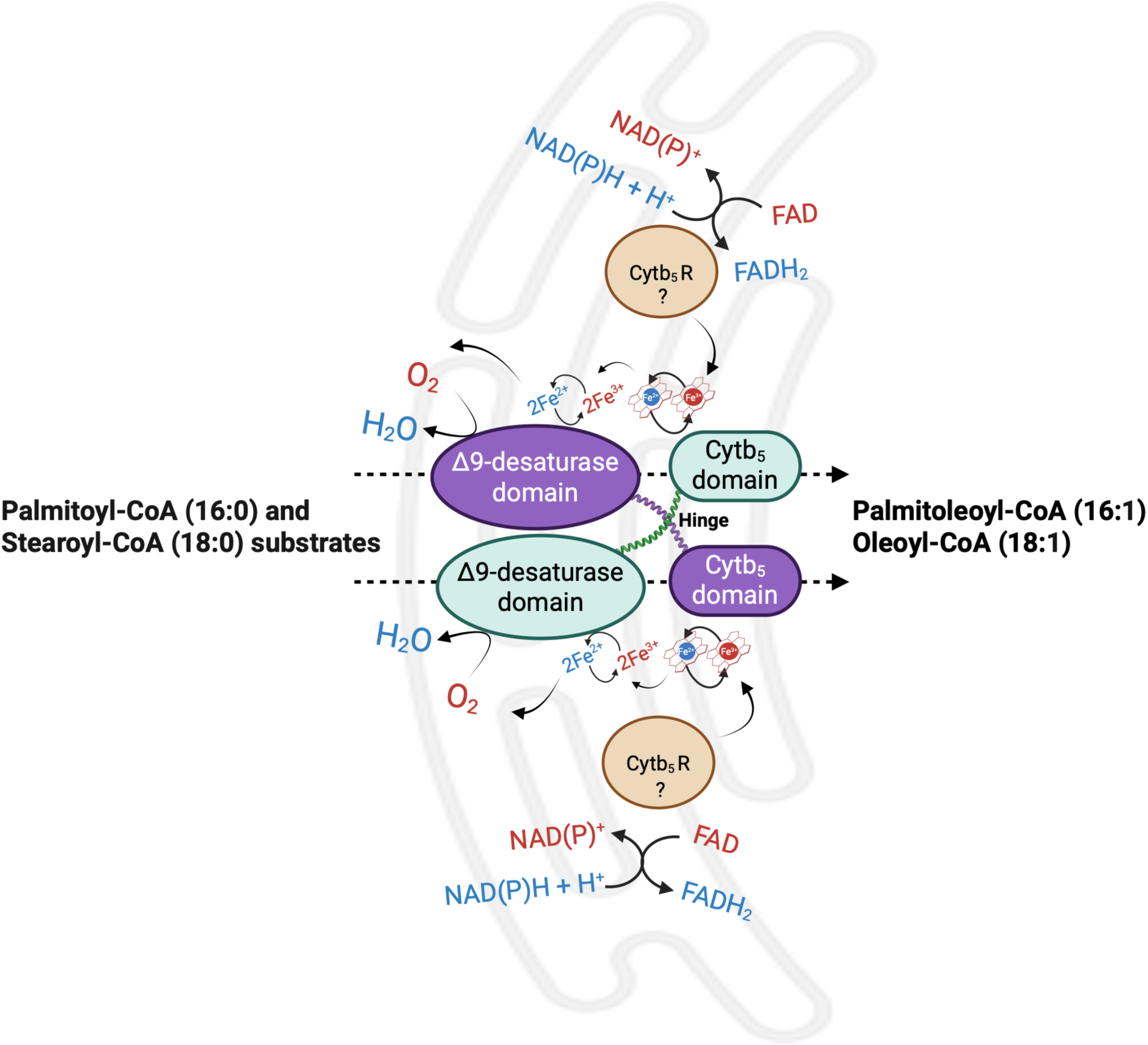
Proposed model for SdeA dimerization and *trans* electron transfer during *A. fumigatus* Δ9-fatty acid desaturation. SdeA is proposed to form a homodimer within the ER membrane through interactions involving the desaturase domain, the fused Cytb5 domain, and the intervening hinge region. Electrons derived from NAD(P)H are proposed to be transferred to an unidentified cytochrome b5 reductase (Cytb5R?), whose flavin cofactor relays reducing equivalents to the heme group of the SdeA Cytb5 domain. The Cytb5 heme is coordinated by His366 and His392. In the proposed dimeric arrangement, the reduced Cytb5 domain of one protomer delivers electrons to the di-iron catalytic center of the neighboring desaturase domain, enabling oxygen activation and double-bond formation in saturated fatty acid substrates. Through this reaction, palmitoyl-CoA and (16:0) and stearoyl-CoA (18:0) substrates are converted into palmitoleic acid (16:1) and oleic acid (18:1), respectively. Structural modeling, biochemical analysis of the Cytb5 module, mutational analysis of the heme-coordinating histidines, and *in vivo* self-association assays support a dimeric architecture compatible with trans electron transfer, although direct interprotomer electron transfer remains to be demonstrated. The hinge region is proposed to maintain the spatial organization required for productive Cytb5-desaturase coupling. This model is consistent with the loss of function observed after deletion of either the Cytb5 domain or the hinge region, despite preservation of ER-associated localization, and with the inability of a detached Cytb5-containing segment to complement the *sdeA*^Δcytb5^ mutant. Reduced and oxidized molecules and iron atoms (Fe^2+^and Fe^3+^) are shown in blue and red, respectively. Filled arrows indicate the proposed direction of electron transfer. Dashed arrows indicate the FA desaturation reaction. Created with BioRender.com.

A central implication of our study is that productive Δ9-desaturation requires spatial coupling between electron transfer and catalysis. Structural studies of mammalian SCD1 have shown how the desaturase core positions a di-iron center and acyl-CoA substrate within the membrane environment for double-bond formation ^18, 20^. In mammals, however, electron delivery requires separate redox partners, Cytb5 and Cytb5R. Recent work showed that mammalian SCD1, Cytb5 and Cytb5R can assemble into stable membrane-organized binary and ternary complexes, and that this organization markedly enhances electron-transfer efficiency ^27^. Thus, efficient desaturation depends not only on redox cofactors but also on the physical organization of the electron-transfer chain.

Our findings suggest that fungal SdeA addresses this same biochemical constraint through a distinct architectural strategy. Rather than relying on a separate membrane-associated Cytb5 partner, SdeA carries a fused C-terminal Cytb5 domain within the same polypeptide chain (Fig. 7). This domain is not dispensable: Cytb5 deletion abolished SdeA function, H366A and H392A substitutions inactivated the enzyme, and MUFA supplementation restored growth, linking these defects directly to impaired Δ9-desaturation. Importantly, the isolated SdeA Cytb5 domain displayed heme-dependent redox responsiveness *in vitro*, yet a detached Cytb5-containing segment failed to restore growth of the *sdeA*^Δcytb5^ strain. Moreover, whereas the truncated SdeA^ΔCytb5^::GFP retained ER-associated localization, the detached Cytb5::mRFP module did not detectably colocalize with SdeA^ΔCytb5^::GFP and accumulated predominantly in punctate cytosolic structures. Together, these data support a model in which the fused Cytb5 domain is not an interchangeable redox appendage but an integral architectural element that must be physically integrated within the native SdeA fusion architecture to enable productive electron delivery to the catalytic center, although targeting and *cis* integration remain formally inseparable here.

The importance of fused Cytb5 domains is not unique to SdeA, but our work places this feature into a mechanistic framework for a pathogenic fungus. A C-terminal Cytb5-like domain was identified decades ago in *S. cerevisiae Ole1* ^21^, and Cytb5-fused desaturases are widespread among fungi and other eukaryotic lipid desaturases ^28^. Similarly, plant and algal Δ6-desaturases contain N-terminal Cytb5 domains required for activity, with conserved histidines mediating heme coordination ^29, 30^. These comparisons suggest that Cytb5 fusion represents a recurrent architectural solution for positioning a heme-containing electron donor near a membrane-embedded desaturase core.

Our data further identify the hinge region as a second critical architectural determinant of SdeA function. The hinge lies between the desaturase core and the Cytb5 domain, precisely where an organizing element would be expected to influence the relative positioning of catalytic and redox modules (Fig. 7). Deletion of the hinge abolished SdeA-dependent growth and virulence, yet did not eliminate ER-associated localization of the protein. Instead, SdeA^Δhinge^::GFP retained reticular fluorescence but also accumulated in large punctate cytosolic structures, consistent with partial destabilization or misassembly. Most importantly, hinge deletion abolished stable SdeA self-association *in vivo*, with only a faint residual signal in one direction. Both baits were efficiently recovered in the direct-capture lanes, indicating loss of interaction rather than failed pulldown; and because the same matrices and conditions were used throughout, the hinge-deleted strain serves as an internal specificity control for the full-length co-IP. Because SdeA^Δhinge^::GFP also accumulated in punctate cytosolic structures, however, we cannot formally distinguish whether the hinge constitutes a direct self-association interface or whether its removal destabilizes SdeA to an extent that precludes assembly. These findings support a model in which the hinge is not a passive linker but a structural organizer required for assembly or stabilization of a catalytically competent SdeA complex. This interpretation parallels the principle illustrated by mammalian SCD1: membrane redox catalysis depends on structural elements that organize electron donors relative to catalytic centers. In the SCD1 system, the transmembrane helices of Cytb5 and Cytb5R stabilize binary and ternary complexes, enhancing electron transfer ^27^. In SdeA, the hinge may serve a conceptually analogous organizational role, not as an equivalent structural element, but by supporting self-association and a dimeric arrangement compatible with *trans* electron transfer, in which the Cytb5 domain of one protomer approaches the catalytic region of the other at a distance within the range over which biological electron tunnelling operates. Although direct demonstration of interprotomer electron transfer remains an important future goal, the combined experimental data support a model in which hinge-dependent self-association contributes to catalytic competence.

Despite the insights provided by our study, the lack of an experimentally resolved structure of full-length fungal SdeA remains an important limitation of the predicted structural model. Our attempts to obtain recombinant full-length 6×His::SdeA or GST::SdeA in bacterial expression systems have so far been unsuccessful (unpublished observations), likely reflecting the intrinsic difficulty of producing a multi-pass membrane desaturase with a fused redox module in a heterologous system. Future work using eukaryotic expression platforms and membrane-mimetic systems will be required to determine whether the predicted *trans* configuration is correct and to elucidate how the hinge, Cytb5 module, and desaturase core are arranged during catalysis. Another unresolved question concerns the upstream redox partner that reduces the Cytb5 domain. By analogy with mammalian and other microsomal desaturase systems, an NAD(P)H-dependent Cytb5R or related flavoprotein is a plausible upstream donor ^18, 27,31^. The importance of Cytb5-dependent redox systems in *A. fumigatus* is further supported by CybE, whose loss impairs hyphal tip growth, membrane organization, and cellular fitness and has been linked to altered sterol metabolism and azole susceptibility^32^. This does not imply that CybE substitutes for the fused Cytb5 domain. Indeed, our data argue that no endogenous Cytb5-containing activity compensated for the removal of the SdeA Cytb5 module. Thus, SdeA represents a specialized architecture in which the redox module is embedded directly within the desaturase polypeptide.

Our analysis of the putative *sdeB* locus further emphasizes the specialization of SdeA in *A. fumigatus*. Although *A. nidulans* encodes two functional Δ9-desaturases, the *A. fumigatus sdeB*-related locus is ambiguously annotated, transcriptionally represented, and lacks a resolved mature coding sequence under the conditions analyzed here. The tagged construct tested did not yield detectable SdeB accumulation, the *sdeB* constructs failed to compensate for *sdeA* depletion during growth or infection, and deletion of the locus did not reveal an evident phenotype or enhance the conditional *xylP::sdeA* defect. These findings caution against assigning Δ9-desaturase function based solely on desaturase-like sequence similarity or locus annotation. Instead, a fused Cytb5 domain, conserved heme-coordinating histidines, an intact desaturase core, and appropriate connecting architecture may define functional SdeA/Ole1-like enzymes in pathogenic fungi and help prioritize candidates for antifungal target discovery.

Taken together, our findings define SdeA as a Cytb5-dependent and hinge-organized Δ9-fatty acid desaturase required for MUFA biosynthesis, growth, and virulence in *A. fumigatus*. The virulence defects observed across SdeA-disrupting alleles further suggest that endogenous MUFA production is required during infection, where environmental lipid availability appears insufficient to compensate for loss of SdeA function. SdeA activity does not reside in the catalytic core alone, but emerges from coordinated assembly of the desaturase domain, fused Cytb5 module, hinge region, and dimeric organization. By exposing architectural features absent from the human Δ9-desaturase - the fused redox-domain interface and hinge-dependent assembly - this architecture provides both a mechanistic model for fungal self-sufficient desaturases and a framework for antifungal strategies targeting lipid desaturation, redox-domain coupling, and membrane adaptation in human fungal pathogens.

## Material and Methods

### Structural modeling of SdeA

SdeA models were built using the AlphaFold3 server (https://alphafoldserver.com/) ^33^. For SdeA (AFUB_091500), two copies of the protein, 2 heme groups, and 4 Fe^3+^ ions were used as input, with a resulting ipTM score of 0.72 and a pTM score of 0.75. Predicted protein domains, including the FA desaturase domain and the Cytb5 domain, were identified using InterProScan/InterPro search. Images were generated using ChimeraX^34^. Predicted aligned error (PAE) plot analysis was performed using the ChimeraX Structure Prediction tab, using AlphaFold3 output. Comparison with human SCD1 crystallographic structure ^18^ (PDB 4ZYO) was performed using the ChimeraX Matchmaker tool.

### Sequence analysis of putative *sdeB* locus

For *sdeB* sequence analyses, the AFUB_090930 sequence from A1163 and the Afu7g05350 sequence from Af293 were retrieved from FungiDB. In addition, partial *sdeB* sequences from the laboratory strains were obtained by Sanger sequencing. The CEA10 *sdeB* sequence was retrieved from the genome assembly deposited at NCBI (GCA_051225625). Genomic sequences were aligned using SnapGene software (version 8.2.2), and chromatograms were inspected to verify the presence or absence of the T200. To validate the functional expression of Afu7g05350, RNA-seq data from 13 *A. fumigatus* experiments were analyzed. The Sequence Read Archive (SRA) accession numbers used were DRR101037, DRR101038, DRR101039, DRR101040, DRR101041, SRR5306457, SRR5878937, SRR5878940, SRR5878944, SRR6163648, SRR6163649, SRR6491493, and SRX9806355. The datasets were identified and accessed via the FungiExpresZ platform ^35^. All SRA datasets were processed using a standardized alignment pipeline. Quality-filtered reads were aligned to the complete reference sequence of the Afu7g05350 locus using Bowtie2 ^36^ with default parameters. The mapped reads files (SAM) were converted to BAM format, sorted, and indexed using SAMtools ^37^. The sorted BAM files were then visualized in the Integrative Genomics Viewer (IGV) ^38^ to manually inspect read coverage and the presence of spliced reads spanning putative intron-exon junctions. This analysis enabled inference of intron positions from RNA-seq alignment gaps, which were compared directly with the current gene annotation for Afu7g05350.

### Strains and growth conditions

The strains used in this study are described in Supplementary Table S1. Strains were maintained in minimal medium [MM; glucose 1% (w/w), 1× high-nitrate salts, 1× trace elements, pH 6.5], as described previously ^39^. When required, pyrithiamine hydrobromide or hygromycin was added at final concentrations of 0.2 µg/ml and 250 µg/ml, respectively. To allow growth of the conditional *xylP::sdeA* mutant, xylose was added to the MM at the concentrations mentioned in each experiment. Conidia were collected with 0.01% Tween 80 solution, filtered through Miracloth, counted with a hemacytometer, and diluted in sterile water or medium to the final concentration required for each assay. When necessary, liquid cultures were grown overnight at 37°C and 180 rpm in shake flasks. For DNA or protein extraction, mycelia were recovered by filtration and stored at -80°C until used.

### Construction of *sdeA* mutants and CRISPR-Cas9 procedures

In all cases, the recovered cassettes constructed here were introduced into the *aft4* genomic Safe haven (Sh) previously described ^25^ using CRISPR-Cas9 methodology optimized for *Aspergillus* ^40, 41, 42^ and the Sh-targeting constructs and PCR validation strategies described previously ^43^. The conditional *xylP::sdeA* mutant and the tagged *sdeA*::GFP and *sdeA*::3×HA ^15^ were used as the recipient strain for most cassette integrations. The primers used in the constructions are listed in Supplementary Table S2. The *sdeA*^Δcytb5^::GFP strain was constructed by deleting the last 378 bp of the *sdeA* gene (AFUB_091500), corresponding to the C-terminal 126 amino acid residues containing the Cytb5 domain from Leu331 to Ala456 (Fig. S1A-B). For this purpose, two DNA fragments were generated from the previously constructed *sdeA*::GFP strain. The first fragment, containing the endogenous *sdeA* promoter region and the truncated *sdeA* coding sequence lacking the Cytb5-encoding region, was amplified using primers IM-315 and IM-751. The second fragment, containing GFP, the *pyrG* marker, and the *sdeA* 3’ regulatory region, was amplified using primers IM-170 and IM-318. These fragments were assembled by *in vivo* recombination in *S. cerevisiae* as previously described ^39^. The resulting cassette was then used as template to generate the donor cassette for CRISPR-Cas9-mediated integration at the Sh locus using primers IM-g48 and IM-g49 (Fig. S1). For the double SdeA-tagged strains, the 3×HA cassette was amplified from the genomic DNA of the *sdeA*::3×HA strain using primers IM-g48 and IM-g49. The resulting PCR product was used as the donor cassette for integration into the Sh locus of the *sdeA*::GFP strain.

The *sdeA*^Δhinge^ strain was generated by deleting 75 bp of the *sdeA* gene, corresponding to the 25-amino-acid hinge region identified in our structural model, spanning Phe298 to Leu322 (Fig. 1 and Fig. S4A-B). To this end, a DNA cassette lacking these 75 bp was generated using wild-type genomic DNA as template. The 5’ fragment, containing the endogenous *sdeA* promoter region and the upstream portion of the *sdeA* coding sequence, was amplified using primers IM-g48 and IM-801. The 3’ fragment, containing the downstream portion of *sdeA* and the *sdeA* terminator region, was amplified using primers IM-802 and IM-g49. The two fragments were fused by PCR using the IM-g48 and IM-g49 primers. The same strategy was applied, using *sdeA::*GFP genomic DNA as the template, to generate the *sdeA*^Δhinge^::GFP cassette. The constructed cassettes were transformed into the *xylP::sdeA* or *sdeA*::3×HA strains. Candidate transformants were screened by diagnostic PCR using primers IM-320 and IM-751, which distinguish the wild-type and Δhinge alleles, and by amplification of the Sh locus using primers IM-803 and IM-804 (Fig. S4C-E). The modified Sh locus was also amplified and sequenced to validate deletion of the 75-bp hinge region of both strains used in this work (Fig. S4F).

A similar strategy was used to generate substitution cassettes containing the *sdeA*^H366A^ and *sdeA*^H392A^ mutations. For the *sdeA*^H366A^ allele, two overlapping fragments were amplified using primer pairs IM-g48 + IM-844 and IM-843 + IM-g49. For the *sdeA*^H392A^ allele, similarly, two overlapping fragments were amplified using primer pairs IM-g48 + IM-846 and IM-845 + IM-g49. The corresponding mutagenic primers introduced the CAC-to-GCC substitutions required to replace His366 or His392 with alanine (Fig. S3A-B). The overlapping fragments were fused by PCR to generate the final Sh donor cassettes. The constructed cassettes were transformed into the *xylP::sdeA* strain, and the modified Sh locus was amplified with primers IM-g48 and IM-g49. The resulting purified fragments were sequenced to validate the relevant His point mutations in the Cytb5 domain (Fig. S3C).

The Cytb5::mRFP strain was generated by ectopically expressing the Cytb5 domain of SdeA under the control of the endogenous *sdeA* promoter. The mRFP::3×HA::ptrA module was amplified from the previously described construct ^15^ using primers IM-815 and IM-448. The CytB5-coding region was amplified from wild-type genomic DNA using primers IM-814 and IM-508, whereas the endogenous *sdeA* promoter region was amplified using primers IM-315 and IM-813. The primers were designed to generate overlapping regions between adjacent fragments, allowing assembly of the complete cassette by overlapping PCR (Fig. S2A). The cassette was inserted into the *xylP::sdeA* strain, and transformants were selected for pyrithiamine resistance. Candidates were screened by PCR using primers IMq-81 and IM-448 (Fig. S2B).

### Construction of the *sdeB* expression, deletion, and complemented strain

To evaluate the ability of *sdeB* to complement the loss of *sdeA*, two ectopic *sdeB* expression cassettes were generated for targeted integration at the Sh locus. The primers used in the constructions are listed in Supplementary Table S2. In these constructs, the *sdeB* sequence corresponding to the Af293 allele (*sdeB*^Af^) was placed under the control of its endogenous regulatory regions or the strong *gpdA* promoter to generate endogenous expression and overexpression cassettes, respectively. Both constructs were transformed into the *xylP::sdeA* strain. For the *sdeB*^Af^ endogenous-expression construct, the cassette was amplified from Af293 genomic DNA using primers IM-g61 and IM-g62, which harbor the homology arms for Sh integration via CRISPR-Cas9 (Fig. S6A). A similar strategy was used to generate the *gpdA*::V5::*sdeB*^Af^ overexpression cassette. The *gpdA* promoter fused to the V5 sequence was amplified from *A. nidulans* genomic DNA using primers IM-g53 and IM-830, whereas the *sdeB*^Af^ sequence and 3’ region were amplified using primers IM-832 and IM-g62. Primer IM-832 contains a 5’ overhang complementary to the V5 sequence, allowing the two fragments to be fused by PCR. The final *gpdA*::V5:: *sdeB*^Af^ cassette was generated by fusion PCR and contained the 5’ and 3’ Sh homology arms required for CRISPR-Cas9-mediated integration at the Sh locus (Fig. S6B). To generate the Δ*sdeB* null mutant, genomic DNA from the wild-type Δ*akuB*^KU80^ strain was used as template to amplify approximately 1.5 kb fragments corresponding to the 5’ (IM-510 + IM-511) and 3’ (IM-513 + IM-514) flanking regions of the *sdeB* locus. The *ptrA* pyrithiamine resistance cassette was amplified using primers IM-512 and IM-329 and used as the selectable marker. The three fragments were assembled by *in vivo* recombination in *S. cerevisiae* ^39^ to generate the Δ*sdeB* deletion cassette (Fig. S6C). The resulting cassette was used to transform the Δ*akuB*^KU80^ and *xylP::sdeA*. Candidates were screened by diagnostic PCR (Fig. S6D) and further confirmed by Southern blot analysis (Fig. S6E). For complementation, the native *sdeB* locus was amplified with the primers IM-639 + IM-640 and cloned into the pBS311 plasmid (a gift from Dr. Robert Cramer), which carries a hygromycin resistance cassette, using NEBuilder HiFi DNA Assembly (New England Biolabs). The resulting construct was transformed into the Δ*sdeB* mutant to generate the complemented strain Δ*sdeB::sdeB*^+^. Complemented transformants were validated by PCR using primers IM-639 and IM-517 (Fig. S6F).

### Southern blot

Southern blot analysis was performed to confirm the correct homologous integration of the *sdeB* deletion cassette at the targeted genomic loci. Genomic DNA from *A. fumigat*us strains was extracted as previously described ^39^. For Southern blotting, *Eco*RI-restricted chromosomal DNA fragments were separated on a 1% agarose gel, transferred to Hybond-N nylon membranes (Cytiva), and processed according to standard procedures^44^. Probe labeling and detection were carried out using the AlkPhos Direct Labeling and Detection System (Cytiva) according to the manufacturer’s instructions. Labeled membranes were exposed to Hyperfilm ECL (Cytiva), and the resulting signals were recorded with the ChemiDoc XRS gel imaging system (Bio-Rad).

### Phenotypic assays

Conidia assays to evaluate the strain’s ability to grow in the presence of glucose or xylose were performed by spotting 1×10^4^ conidia on 90 mm agar plates. The plates were incubated for the periods described in each experiment at the indicated temperature.

### Cloning of the Cytb5 fragment, recombinant expression, and purification

The coding sequence of the cytb5 region spanning Asp323 to Ala456 was amplified from the plasmid pET15b, which contained the entire *E. coli* codon-optimized *sdeA* coding sequence (Biobasic), using the primers IM-858 and IM-850 (Table S2). These sequences were subcloned into the pET15b plasmid using NEBuilder HiFi DNA Assembly (New England Biolabs) methodology to create pET15b-cytb5. *E. coli* DH10B was transformed with the constructs for propagation of the plasmids. Plasmid extractions were performed, and *E. coli* BL21Rosetta was transformed with the constructs for protein expression. Single colonies of Rosetta cells harboring each plasmid were cultured in LB medium containing carbenicillin (100 μg/ml) and grown for 16 hours at 37°C in an orbital shaker (250 rpm). The culture was then diluted into 1 L of LB medium containing 100 μg/ml carbenicillin and incubated at 37°C until the OD_600_ reached 0.5. IPTG was then added to a final concentration of 0.4 mM, and the culture was incubated at 18°C overnight. Cells were harvested by centrifugation, and the pellet was washed and suspended in lysis buffer (50 mM Tris-HCl, pH 7.5, 250 mM NaCl, 1% Triton X-100, 10 mM imidazole, 30 μg/mL lysozyme, 1 U/mL Benzonase, 1 mM PMSF, and 1 mM DTT). Cell lysis was performed by sonication, and the crude extract was centrifuged at 15,000 rpm for 45 min at 4°C to remove insoluble material. The resulting supernatant was collected, filtered, and used for protein purification. Recombinant proteins were purified by immobilized metal affinity chromatography (IMAC) using a HisTrap column (Cytiva). Purification conditions were optimized by applying an imidazole gradient according to the manufacturer’s recommendations. Following purification, imidazole was removed by dialysis against buffer containing 50 mM Tris-HCl (pH 7.5), 250 mM NaCl, and 5 mM β-mercaptoethanol. Recombinant protein concentrations were determined spectrophotometrically at 280 nm. The extinction coefficient for reduced Cytb5 proteins was determined using the ProtParam tool (http://www.expasy.ch/tools/protparam.html).

### UV-visible spectroscopy of recombinant 6×His::Cytb5

UV-visible absorption spectra of purified recombinant 6×His::Cytb5 were obtained to assess heme incorporation into the protein, as previously described ^24^. Purified protein (10 µM) in buffer containing 50 mM Tris-HCl (pH 7.5), 250 mM NaCl, and 5 mM β-mercaptoethanol was analyzed in a spectrophotometer with quartz cuvettes. Spectra were collected between 350 and 700 nm at 25°C, with the relevant absorption ranges shown in the respective figures. The oxidized spectrum was recorded directly from the purified protein sample. For reduction, sodium dithionite was added to a final concentration of 5 mM, and the spectrum was reacquired immediately after mixing. Buffer alone was used for blank subtraction. The position of the Soret peak and the visible absorption bands were used to assess the redox-responsive properties of the heme-bound Cytb5 module.

### *A. fumigatus* protein extraction, co-immunoprecipitation (Co-IP), and Western blot

Total protein extracts from the relevant tagged strains were prepared using B250 lysis buffer ^45^, according to previously described extraction procedures ^46^, and quantified using Pierce 660 nm Protein Assay (Thermo-Fisher). For co-immunoprecipitation assays, 5 mg of crude protein extract were incubated with 10 µL of EZview Red Anti-HA Affinity Gel (Sigma) or 10 µL of ChromoTek GFP-Trap agarose to immunoprecipitate 3×HA- or GFP-tagged proteins, respectively. Cell extracts and affinity resins were incubated with gentle mixing for 2 h at 4 °C. After incubation, the resins were washed four times with B250 buffer. Bound proteins were eluted in Laemmli sample buffer and analyzed by 12% SDS-PAGE. Proteins were then transferred to PVDF membranes and detected by Western blot using α-HA (Sigma) or α-GFP (Sigma) antibodies, as indicated for each experiment. Primary antibodies were incubated overnight at 4 °C in TBST containing 3% skimmed milk, followed by incubation with horseradish peroxidase (HRP)-conjugated secondary IgG antibody in TBST for 2 h at room temperature. Chemiluminescent detection was performed using Clarity Western ECL Substrate (Bio-Rad). Images were acquired with the ChemiDoc XRS gel imaging system.

### Confocal fluorescence microscopy

Confocal fluorescence microscopy was performed using a Zeiss LSM 900 laser-scanning confocal microscope mounted on an Axio Observer Z1/7 inverted microscope and equipped with an Airyscan 2 detector and a Plan-Apochromat 63×/1.40 oil immersion objective. Nuclei were stained with Hoechst 33342 (10 μg/mL) for 20 min at 37 °C prior to imaging. GFP, mRFP, and Hoechst 33342 fluorescence signals were excited using the 488, 561, and 405 nm laser lines, respectively. Emission was collected at 490-575 nm for GFP, 565-700 nm for mRFP, and 400-495 nm for Hoechst 33342. Images were acquired in Airyscan Super Resolution mode (2D, Auto) and processed using ZEN 3.11 software (Zeiss). For comparative analyses, images within each experiment were acquired and processed using identical microscope settings

### Virulence assay in the *Galleria mellonella* model

Survival assays in *G. mellonella* were performed as previously described ^47^. Larvae were maintained in sterile glass flasks fitted with modified lids containing a central opening covered with ultra-fine stainless-steel mesh to allow ventilation and fed as described ^48^. For each experimental group, 15 healthy larvae of similar weight (approximately 350-380 mg) were selected. Each larva was inoculated with 10 µL of a conidial suspension (1×10^8^ conidia/mL) of the wild-type or mutant *A. fumigatus* strains by injection into the hemocoel through the last left proleg. Larvae injected with PBS were used as a negative control. After inoculation, larvae were incubated at 37°C in the dark, and survival was monitored daily for 15 days. Larvae that failed to respond to touch stimulation were scored as dead. Each infection experiment was performed three independent times, using fifteen larvae per group per experiment. Survival curves were analyzed using the log-rank (Mantel-Cox) test in GraphPad Prism 10, and differences were considered significant at P < 0.05. Significant differences were determined as described in the corresponding figure legends.

### Statistics and reproducibility

All statistical analyses were performed using GraphPad Prism 10. The statistical tests used for each experiment are indicated in the corresponding figure legends. Unless otherwise noted, all experiments were performed with a minimum of three biologically independent samples. For qualitative growth assays, microscopy images, immunoblots, and co-immunoprecipitation assays, representative results from independently repeated experiments are shown whenever appropriate. Statistical significance was set at P < 0.05. Sample sizes and number of replicates are indicated in each figure legend.

## Acknowledgments

We are grateful to Dr. Gustavo Henrique Goldman for providing the *A. fumigatus* A1163 and CEA17 strains, Dr. Robert Cramer for providing the pBS311 plasmid, and Dr. Jéssica Hilário Bonomo for technical assistance in the laboratory. We also acknowledge the National Institute of Science and Technology in Fungal Virulence (INCT FUNVIR) for scientific support.

## Author Contributions

I.M. and J.E.M.C. conceptualized and designed experiments. J.E.M.C., G.Y.F., J.H.T.M.F., and N.F.B. performed experiments and data analysis under the supervision and guidance of I.M., A.F.C., and L.T.R. L.T.R. and N.F.B. were responsible for protein modeling and structural analysis. E.D. collected and analyzed *sdeA* and *sdeB* DNA sequences. P.A.B. helped with laser-scanning confocal microscopy and analysis. I.M., A.F.C., and L.T.R. provided financial support. I.M. and J.E.M.C. wrote the original manuscript. I.M. coordinated all the work. All authors reviewed, edited, and approved the manuscript.

## Data availability

The AlphaFold3-predicted SdeA structural model generated and analyzed during the current study and depicted in Fig. 1 has been deposited in ModelArchive and is publicly available at https://www.modelarchive.org/doi/10.5452/ma-vy3qn. The fungal DNA and protein sequences analyzed in this study correspond to publicly available sequences deposited in FungiDB, including AFUB_091500, AFUB_090930, and Afu7g05350. Publicly available RNA-seq datasets used for transcript-alignment inspection of the *sdeB*locus are cited in the manuscript and were not generated in this study. All other data supporting the findings of this study are available from the corresponding author on reasonable request.

