## Supplementary Information for "The self-sufficient *Aspergillus fumigatus* Δ9-fatty acid desaturase SdeA requires a fused cytochrome b5 domain and hinge-dependent self-association for virulence"

To whom correspondence should be addressed:

**This file includes:**

Supplementary Table 1

Supplementary Table 2

Supplementary Figs. 1-12

Supplementary References

**Supplementary Table 1:** *A. fumigatus* strains used in this study.

| Name | Genotype | Reference |
|---|---|---|
| ΔKU80 pyrG1 | Δ*akuB; pyrG- MAT1-1* | s |
| *xylP::sdeA* | *pyrG*::*xylP::sdeA*; Δ*akuB* | [2] |
| *sdeA*::3×HA | *sdeA::*3×HA::*prtA;* Δ*akuB* - $Ptr^R$ | [2] |
| *sdeA*::GFP | *sdeA*::GFP::*pyrG;* Δ*akuB* | [2] |
| *sdeA*::GFP *sdeA*::3×HA (Sh) | *sdeA*::GFP::*pyrG;* Δ*akuB; sdeA*::3×HA::*ptrA* (Sh) - $Ptr^R$ | This study |
| *sdeA*::3×HA $sdeA^{\Delta hinge}$::GFP (Sh) | *sdeA*::3×HA::*ptrA;* Δ*akuB;* $sdeA^{\Delta hinge}$::GFP::*pyrG;* (Sh) - $Ptr^R$ | This study |
| $sdeA^{\Delta cytb5}$::GFP (Sh) | *pyrG*::*xylP::sdeA*; Δ*akuB;* $sdeA^{\Delta cytb5}$::GFP (Sh) | This study |
| $sdeA^{\Delta cytb5}$::GFP (Sh) cytB5:: mRFP (CytB5+) | *pyrG*::*xylP::sdeA*; Δ*akuB;* $sdeA^{\Delta cytb5}$::GFP (Sh); cytB5::mRFP::3×HA::*prtA* - $Ptr^R$ | This study |
| $sdeA^{\Delta hinge}$ (Sh) | *pyrG*::*xylP::sdeA*; Δ*akuB;* $sdeA^{\Delta hinge}$ (Sh) | This study |
| $sdeA^{H366A}$ | *pyrG*::*xylP::sdeA*; Δ*akuB;* $sdeA^{\Delta H366A}$ (Sh) | This study |
| $sdeA^{H392A}$ | *pyrG*::*xylP::sdeA*; Δ*akuB;* $sdeA^{\Delta H392A}$ (Sh) | This study |
| Δ*sdeB* | Δ*sdeB::ptrA;* Δ*akuB* | This study |
| Δ*sdeB::*$sdeB^+$ | Δ*sdeB::sdeB;* Δ*akuB* - $Ptr^R$, $Hyg^R$ | This study |
| *xylP::sdeA;* Δ*sdeB* | *pyrG*::*xylP::sdeA*; Δ*sdeB::ptrA;* Δ*akuB* | This study |
| *xylP::sdeA;* $sdeB^{Af293}$ | *pyrG*::*xylP::sdeA*; Δ*akuB;* $sdeB^{Af293}$ (Sh) | This study |
| *xylP::sdeA; gpdA::V5::*$sdeB^{Af293}$ | *pyrG*::*xylP::sdeA*; Δ*akuB; gpdA::V5::*$sdeB^{Af293}$ (Sh) | This study |

Sh: *aft4* Safe haven locus described in Furukawa, van Rhijn [3].

**Supplementary Table 2:** Primers used in this study.

| Primer name* | Sequence (5'⟶ 3') |
|---|---|
| IM-170 | GGAACACGGGGAATGAGTAAAGGAGAAGAACTTTTCA |
| IM-174 | TGATCTGGGTATCTTGAAAAGC |
| IM-315 | GTAACGCCAGGGTTTTCCCAGTCACGACGCGCAGAGCCTTATCACGTACCC |
| IM-318 | GCGGATAACAATTTCACACAGGAAACAGCAGAGGAGGAGCCACGGTGACAA |
| IM-319 | GCCTCTGATAGCTATATGCC |
| IM-320 | GTAACGCCAGGGTTTTCCCAGTCACGACGCCGTTACACCGACACTGACAAG |
| IM-329 | CTATCATGGGGTGACGATGAGCCG |
| IM-391 | AGCTACATTTATCTTCAATG |
| IM-392 | AGCCTGAAGTCCTGACCATC |
| IM-448 | CTATCATGGGGTGACGATGAGCCG |
| IM-508 | ACCACCGCTACCACCTCCTGCCGCATCAGCGGTGGGAA |
| IM-510 | GTAACGCCAGGGTTTTCCCAGTCACGACGGTTTTAGAGCAATTCGCCCT |
| IM-511 | TCAATTGCCCGTCTGTCAGATCTGTGTTCGTCGCAGAAGTTA |
| IM-512 | GATCTGACAGACGGGCAATTGA |
| IM-513 | CGGCTCATCGTCACCCCATGATAGTAGATTGGGACGAGTTTGTT |
| IM-514 | GCGGATAACAATTTCACACAGGAAACAGCTCTAAGGTACCTAAGGAGTC |
| IM-515 | GATTCTGAACGTTGAAAGCG |
| IM-639 | GCCAGATCTACGCGTTTAATCAAGCTGCGGCGAGTCCT |
| IM-640 | CGGCCGCCACCGCGGTTAATATAGACGATGACGTATCGGGGATC |
| IM-735 | GGCATAGTCGGGAACGTCATAAG |
| IM-745 | CGTCCAACATCAGCGTCATG |
| IM-746 | TCCGTTCCATTACGCCTTCC |
| IM-751 | AAAGTTCTTCTCCTTTACTCATTCCCCGTGTTCCCTGGTCGAGAGGGATACCCC |
| IM-756 | TTTGTATAGTTCATCCATGC |
| IM-769 | TTACACATGGCATGGATGAACTATACAAATAAGCATTTCCTTACACTATATCAC |
| IM-801 | GGGAAGCTGGTCGAGAGGGATACCCCAATCCTGCTTCAGATCATAGGCTAGACC |
| IM-802 | AGCTTGGTCTAGCCTATGATCTGAAGCAGGATTGGGGTATCCCTCTCGAC |

| | |
|---|---|
| IM-803 | GGGTCTCCAACATGTCATTAGC |
| IM-804 | GCAAAGACATGGTCGAGTTCGT |
| IM-813 | AGAGGGATACCGTCGATCTTCTTCTGCATGTTGCCGACCTTTTGCGTTTCT |
| IM-814 | ATGCAGAAGAAGATCGACGGTATCCCTCTCGACCAGC |
| IM-815 | GCGGCAGGAGGTGGTAGCGGTGGTATGGCCTCCTCCGAGGAC |
| IM-830 | AGGGTTAGGGATAGGCTTACCCATTGTGATGTCTGCTCAAGCGGGGTA |
| IM-832 | ATGGGTAAGCCTATCCCTAACCCTCTCCTCGGTCTCGATTCTACGGGAGCCGGGCGCGCCATGATCTCTGATGACGTCCAGGG |
| IM-843 | GCTGGTGTTGTTCACGATGTGACCGATTTCATCAAGGACGCCCCTGGTGGCAAAGCTATG |
| IM-844 | GCCAATGCCGGAGTTGATCATAGCTTTGCCACCAGGGGCGTCCTTGATGAAATCGGTCAC |
| IM-845 | AGGATGCTACTGCCATGTTCAACGGTGGAGTTTACAACGCCTCAAATGCTGCCCACAACC |
| IM-846 | CGCATGGTAGACAGCAGGTTGTGGGCAGCATTTGAGGCGTTGTAAACTCCACCGTTGAAC |
| IM-849 | CAGCAGCGGCCTGGTGCCGCGCGGCAGCCATATGCTCGAGTTCCGTGCCAATGAAATCG |
| IM-850 | CCAACTCAGCTTCCTTTCGGGCTTTGTTAGCAGCCGGATCTCATGCCGCATCAGCGGTG |
| IM-858 | CAGCAGCGGCCTGGTGCCGCGCGGCAGCCATATGCTCGAGGATTGGGGTATCCCTCTCGACC |
| IM-q81 | CCAGCTTCCCGTCATGGA |
| IM-g48 | GCAGGAGCAAAAACAGGCCGGGAAGATATTGCCTAGGAGTTCTGCGCTGGGCGCAGAGCCTTATCACGTACCC |
| IM-g49 | CTGACAGTTCATGGAATAGTAGAATGGTATGATACATTACGTAACCGACCAGCTACATTTATCTTCAATG |
| IM-g53 | GCAGGAGCAAAAACAGGCCGGGAAGATATTGCCTAGGAGTTCTGCGCTGGGCGTTGGTTGAATTTAGAACGTGG |
| IM-g61 | GCAGGAGCAAAAACAGGCCGGGAAGATATTGCCTAGGAGTTCTGCGCTGGCATGACGGGGAGTGTTGGC |
| IM-g62 | CTGACAGTTCATGGAATAGTAGAATGGTATGATACATTACGTAACCGACCGGATCTGCAGAATACTATGCCAC |

* For primers location, refer to Figures S1, S2, S3, S4 and S6. Primers named as IM-g were used in the CRISPR-Cas9 procedures for cassette integration at the Safe haven (Sh) locus following the protocol described in Campanella, Arentshorst [4].

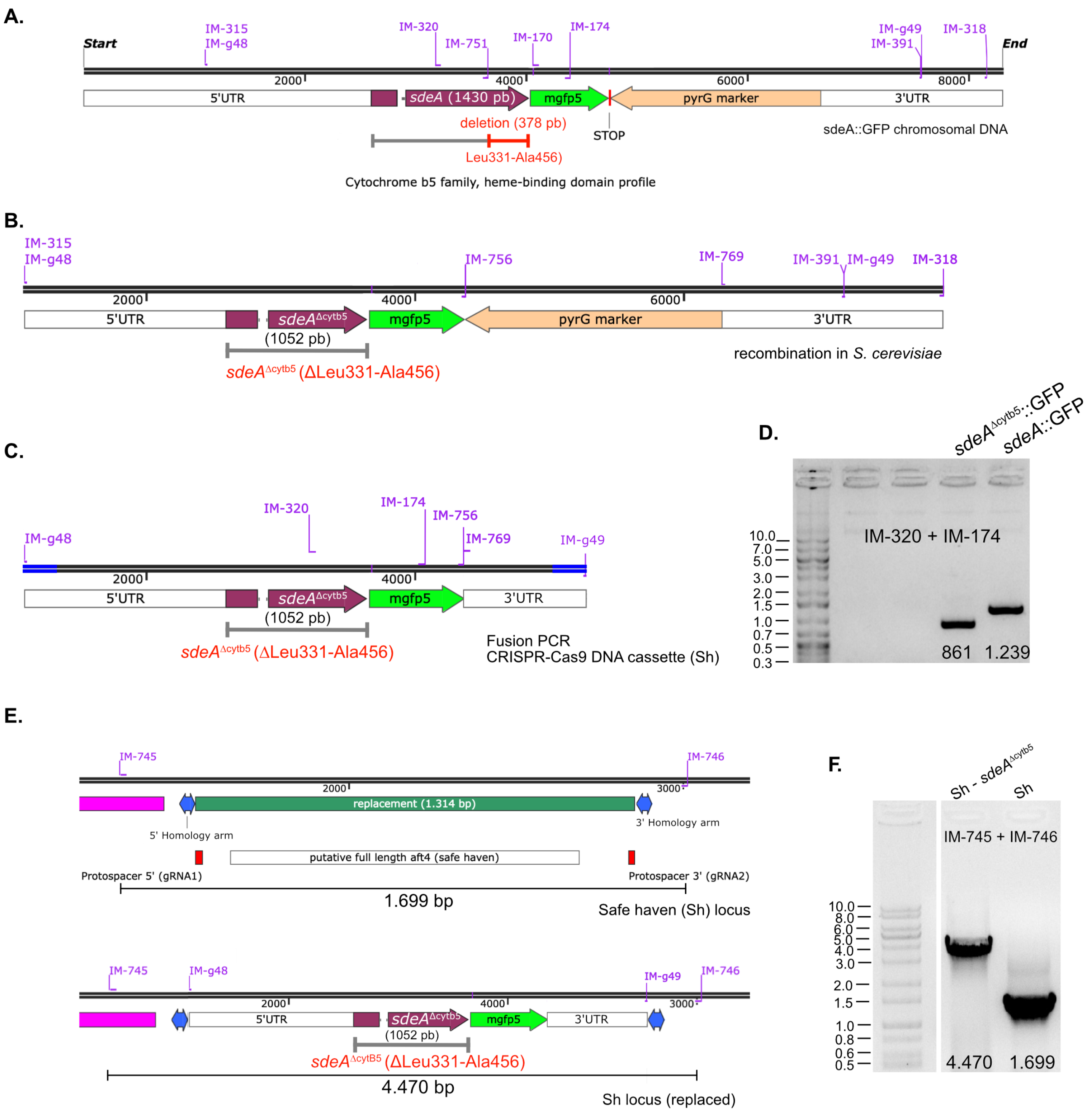


**Figure S1. Generation and validation of the *sdeA*$^{\Delta cytb5}$::GFP mutant strain.** (A-B) Strategy used to generate the *sdeA*$^{\Delta cytb5}$::GFP construct. Primers IM-315 and IM-751 were used to amplify the *sdeA*$^{\Delta cytb5}$::GFP fragment, whereas primers IM-170 and IM-318 amplified the GFP::pyrG::3'UTR fragment from the parental *sdeA::*GFP strain. Both fragments were recombined *in vivo* in *Saccharomyces cerevisiae* to generate the *sdeA*$^{\Delta cytb5}$::GFP::*pyrG* substitution cassette. The deleted region corresponds to 378 bp encompassing residues Leu331 to Ala456 (red line), which include the predicted cytochrome b5 heme-binding domain (B). (C) Assembly of the marker-free donor cassette used for CRISPR-Cas9-

mediated insertion of *sdeA*$^{\Delta cytb5}$::GFP into the Safe haven (Sh) locus in the conditional mutant *xylP::sdeA*. A repair cassette targeting the Sh locus was generated from the substitution cassette shown in B using primer pairs IM-g48+IM-756 and IM-769+IM-g49. The resulting fragments were fused by PCR with primers IM-g48+IM-g49, generating a final donor cassette flanked by 50-bp homology arms corresponding to the Sh locus. (D) PCR-based validation of a positive transformant carrying the *sdeA*$^{\Delta cytb5}$::GFP cassette. Amplification with primers IM-320 and IM-174 yielded the expected 861 bp fragment spanning part of *sdeA* lacking the Cytb5-coding region and the 5' portion of GFP. As a control, the corresponding fragment amplified from the wild-type *sdeA*::GFP strain yielded a 1,239 bp fragment. (E) Schematic representation of CRISPR-Cas9-mediated integration of *sdeA*$^{\Delta cytb5}$::GFP cassette into the Sh locus. The donor construct replaced the endogenous Sh region through homology-directed repair. (F) Validation of correct donor cassette integration at the Sh locus using distal primers IM-745 and IM-746. The mutant strain yielded the expected 4,470-bp amplicon, whereas the wild-type Sh locus generated a 1,699-bp fragment.

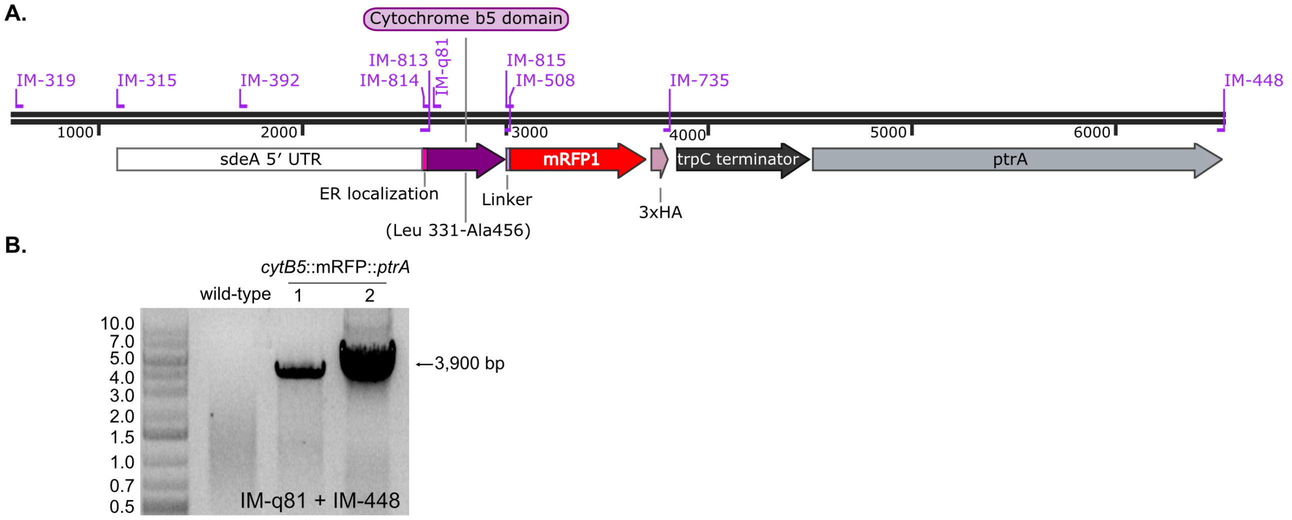


**Figure S2. Construction and validation of the C-terminal Cytb5-containing segment fused to mRFP for *trans*-complementation assays.** (A) Schematic representation of the ectopic expression cassette used to assess whether a detached Cytb5-containing C-terminal segment can functionally complement the truncated *sdeA*$^{\Delta cytb5}$::GFP allele *in trans*. To assemble the construct, the *sdeA* promoter region carrying a sequence overlap with the 5′ end of *cytb5* was amplified with primers IM-319 and IM-813. The coding region corresponding to the Cytb5-containing segment, including an initiation codon, was amplified with primers IM-814 and IM-508. Primer IM-508 was designed with homology to the 5′ end of a previously generated mRFP::3×HA::*trpC*::*ptrA* cassette [5], allowing fusion to the fluorescent module. The mRFP::3×HA::*trpC*::*ptrA* fragment was amplified with primers IM-815 and IM-448. Purified fragments were fused by PCR using primers IM-315 and IM-448, generating the final *cytb5*::mRFP::3×HA::*ptrA* cassette. A GGGSGG linker was introduced between *cytb5* and mRFP. To preserve subcellular targeting of the detached module, the construct retained the QKKID sequence from the upstream hinge-derived region, previously proposed to act as a endoplasmic reticulum (ER) localization determinant [5]. This design was intended to minimize the possibility that a lack of trans-complementation would result from defective ER targeting rather than from the inability of the isolated Cytb5-containing segment to support SdeA function. (B) PCR-based validation of positive transformants carrying the ectopically integrated *cytb5*::mRFP::3×HA::*ptrA* cassette. The construct was introduced into the *xylP::sdeA*; *sdeA*$^{\Delta cytb5}$::GFP (Sh) recipient strain, and transformants were selected by pyrithiamine resistance. Correct candidates (1 and 2) were confirmed by PCR for the presence of the specific 3,900 bp fragment.

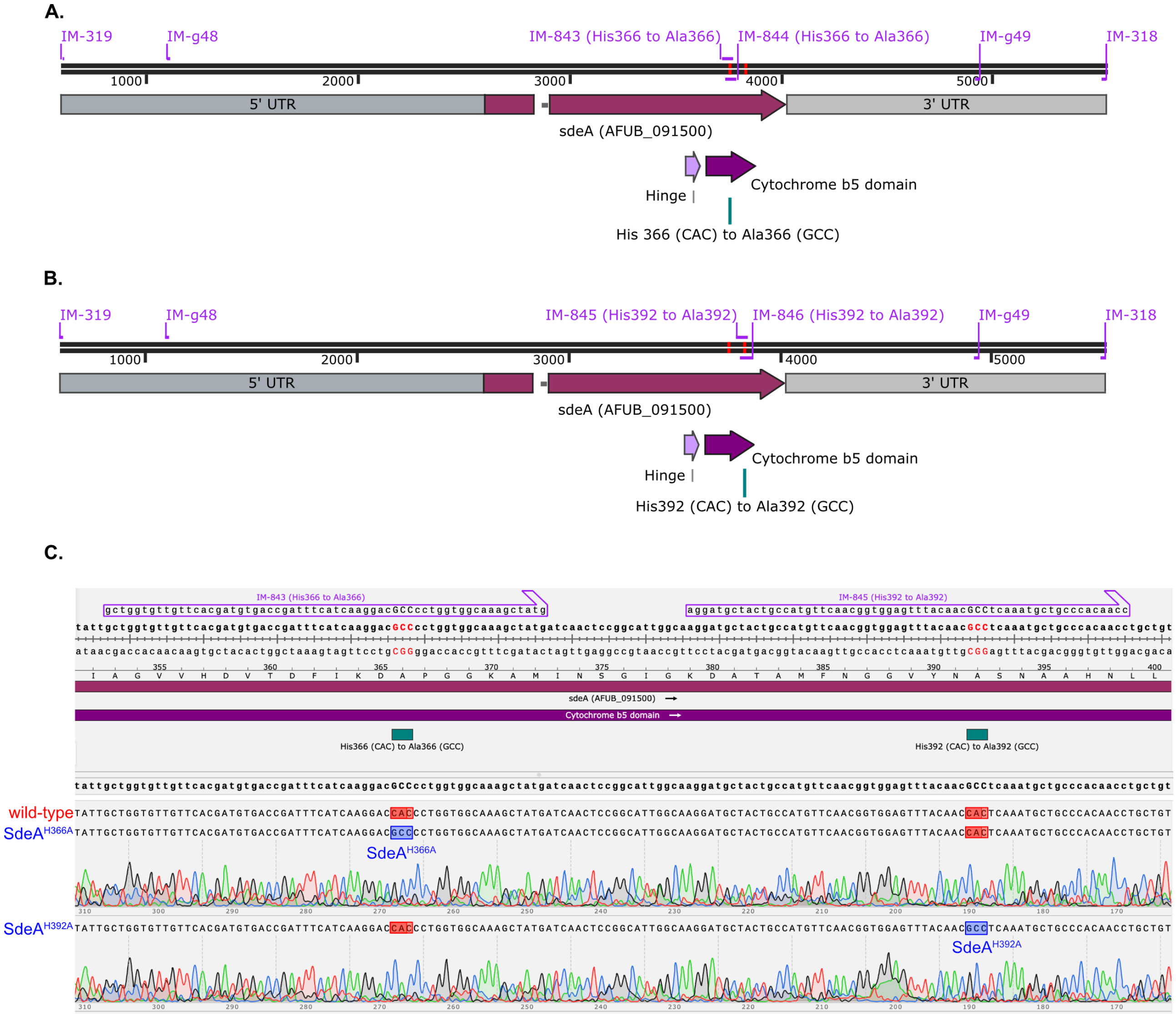


**Figure S3. Construction and sequencing validation of full-length SdeA Cytb5 histidine mutants.** (A) Strategy used to generate the full-length *sdeA*$^{H366A}$ mutant allele. Two overlapping fragments were amplified by PCR using mutagenic primers (IM-843 and IM-844) carrying the H366A substitution within the Cytb5 domain. The 5′ fragment, spanning the endogenous promoter and the 5′ portion of the *sdeA* open reading frame, was amplified with primers IM-319 and IM-844. The 3′ fragment, containing the overlapping mutated region and the 3′ portion of *sdeA* including its endogenous terminator, was amplified with primers IM-843 and IM-318. The H366 residue, predicted to coordinate the heme prosthetic group, was replaced from CAC to GCC. (B), Strategy used to generate the full-length *sdeA*$^{H392A}$ mutant allele. As in A, two overlapping PCR fragments were amplified using mutagenic primers carrying the H392A substitution. The 5′ fragment was generated with primers IM-319 and IM-846, whereas the 3′ fragment was amplified with primers IM-845 and IM-318. The H392 residue within the Cytb5 domain was similarly replaced from CAC to GCC. For both mutant alleles, the purified overlapping fragments were fused by PCR using primers IM-g48 and IM-g49, generating full-length donor cassettes of approximately 3.84 kb flanked by 50-bp homology arms corresponding to the *aft4* Sh locus. The resulting *sdeA*$^{H366A}$ and *sdeA*$^{H392A}$ alleles, containing the endogenous promoter, mutated ORF, and endogenous terminator, were introduced into the conditional *xylP::sdeA* strain by CRISPR-Cas9-mediated integration at the Sh locus.

(C) Sequencing validation of the mutant alleles integrated at the Sh locus. Alignment and representative chromatograms confirmed the expected point substitutions in positive transformants, showing conversion of His366 (CAC) to Ala366 (GCC) in the *sdeA*$^{H366A}$ strain and His392 (CAC) to Ala392 (GCC) in the *sdeA*$^{H392A}$ strain.

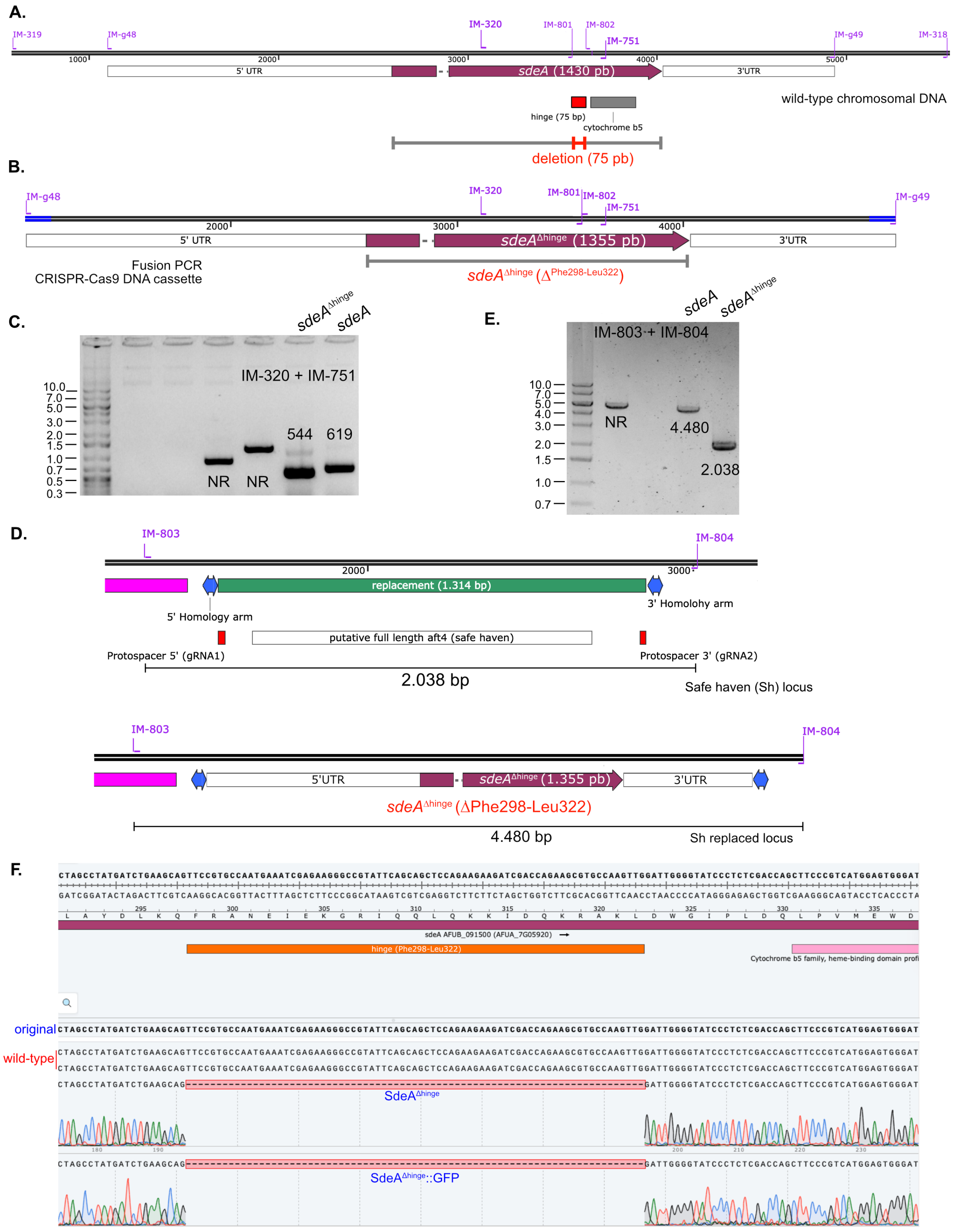


**Figure S4. Construction and validation of the *sdeA*$^{\Delta hinge}$ mutant strain.** (A) Schematic representation of the wild-type *sdeA* locus highlighting the 75-bp region encoding the 25-

residue hinge segment located immediately upstream of the Cytb5 domain identified by the AlphaFold 3 model. Primer positions used for mutant construction and validation are indicated. (B) Strategy used to generate the *sdeA*$^{\Delta hinge}$ donor cassette for CRISPR-Cas9-mediated integration. The hinge-coding segment was removed by overlapping PCR using the primers IM-319 + IM-801 and IM-802 + IM-318 to generate the independent fragments. The external primers IM-g48 and IM-g49 were used in the fusion PCR to generate the final donor cassette. This product contains the truncated *sdeA*$^{\Delta hinge}$ allele and 50-bp terminal homology regions corresponding to the *aft4* Sh locus, enabling targeted integration in the conditional *xylP::sdeA* background. (C) PCR-based validation of positive *sdeA*$^{\Delta hinge}$ transformants using primers IM-320 and IM-751. Amplification of the mutant allele yielded the expected shorter fragment relative to the wild-type *sdeA* allele (544 bp versus 619 bp for the wild-type *sdeA* allele), consistent with deletion of the 75-bp hinge-coding region. (D) Schematic representation of CRISPR-Cas9-mediated integration of the *sdeA*$^{\Delta hinge}$ cassette into the *aft4* Sh locus. (E) Validation of correct donor cassette integration at the Sh locus using distal primers IM-803 and IM-804. The mutant strain yielded the expected 4,480 bp amplicon, whereas the wild-type Sh locus generated a 2,038 bp fragment. (F) Sequencing validation of the *sdeA*$^{\Delta hinge}$ allele. Alignment and representative chromatograms confirmed the precise deletion of the 75-bp segment (Phe298-Leu322) corresponding to the hinge region in independent positive transformants.

A.

B.

C.

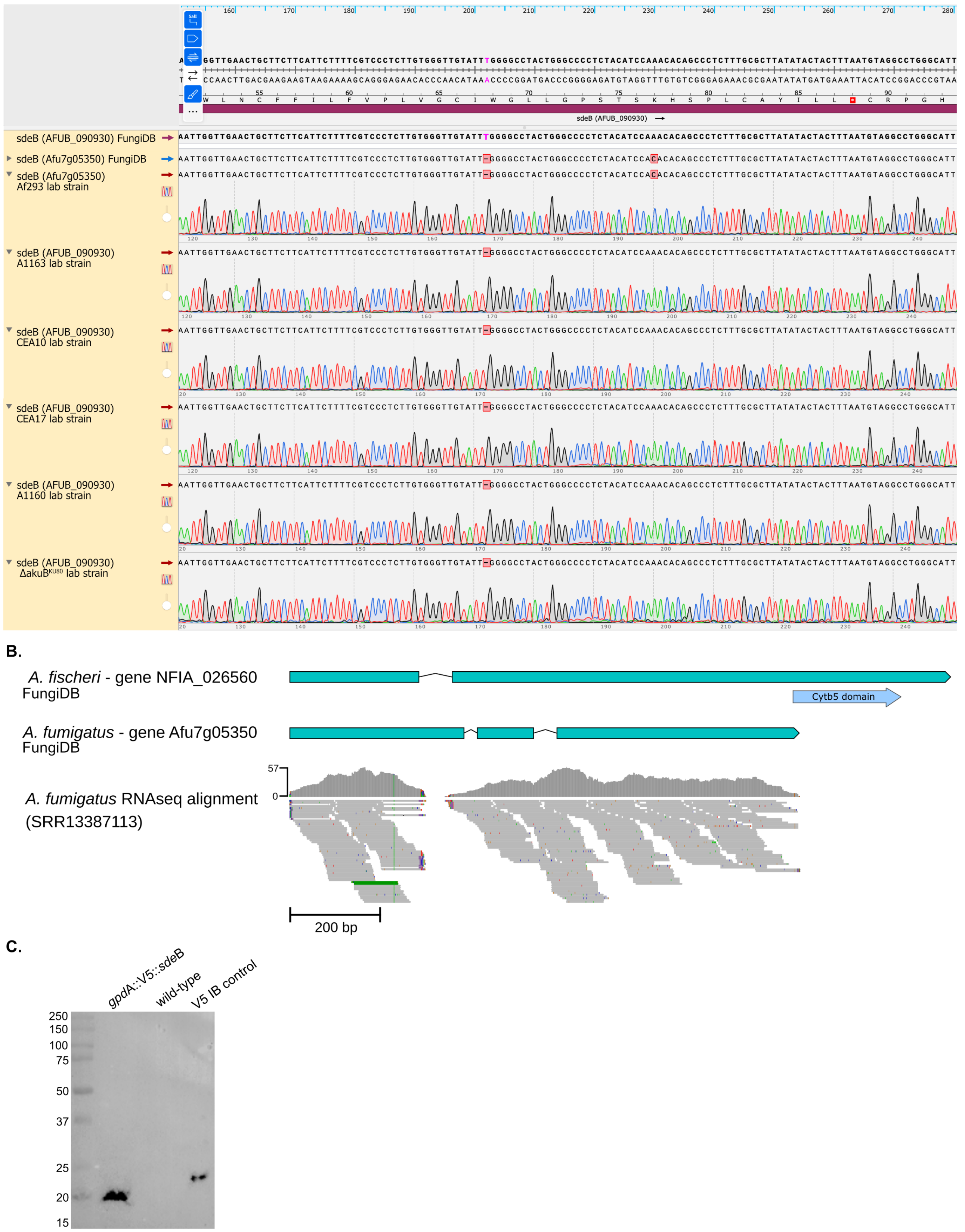


**Figure S5. Sequence, annotation and protein-expression analysis of the putative *sdeB* locus.** (A) Sanger sequence analysis of the *sdeB* locus in the relevant *A. fumigatus*

strain backgrounds. The deposited A1163 AFUB_090930, annotated as a pseudogene in FungiDB (purple arrow) contains a thymine insertion at position 200 (T200), which shifts the reading frame and generates a downstream premature stop codon (asterisk). By contrast, the Af293 Afu7g05350 sequence deposited in FungiDB (blue arrow) and the *sdeB* sequences amplified from the laboratory strains analyzed here (red arrows) do not contain this insertion, preserving the predicted *sdeB* reading frame. (B) Comparison of the annotated *sdeB*-related gene models in *A. fischeri* NFIA_026560 and *A. fumigatus* Afu7g05350 with RNA-seq read alignments mapped to the *A. fumigatus sdeB* genomic locus. The *A. fischeri* ortholog contains a predicted C-terminal Cytb5 domain, whereas the currently annotated *A. fumigatus* Afu7g05350 model lacks the corresponding C-terminal region. The lower panel shows representative RNA-seq reads (SRR13387113) aligned to the *A. fumigatus* locus, revealing splice junctions and intron-containing regions that do not fully match the current Afu7g05350 annotation. Scale bar, 200 bp. (C) Western blot analysis of the ectopic *gpdA*::V5::*sdeB*$^{Af}$ construct using an anti-V5 antibody. A low-molecular-weight V5-reactive species of approximately 20 kDa was detected in the *gpdA*::V5::*sdeB*$^{Af}$ strain, whereas no V5-reactive band corresponding to the predicted intact SdeB$^{Af}$ protein (~40 kDa) was observed under the conditions tested. The wild-type extract was used as a negative control, and a V5-tagged protein (V5::Rho1) extract was included as an immunoblot (IB) control.

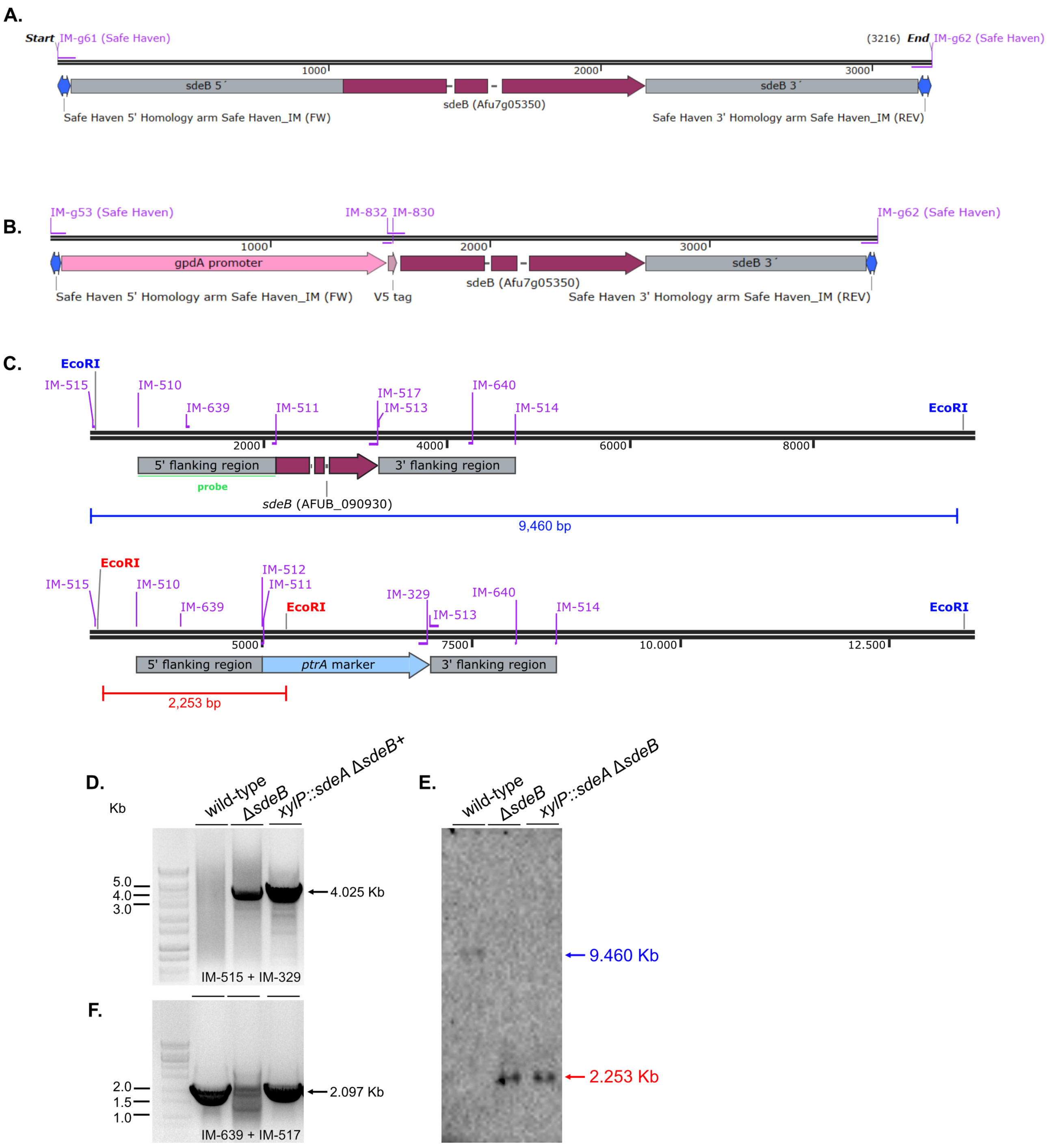


**Figure S6. Construction and validation of the *sdeB* mutants.** (A) Strategy used to generate the ectopic *sdeB* Sh cassette under the control of its endogenous regulatory regions. The construct contains the 5' *sdeB* region, the *sdeB*$^{Af}$ coding sequence (Afu7g05350), and the 3' *sdeB* region. Primers used for amplification and assembly are indicated in the scheme. (B) Strategy used to generate the ectopic *gpdA*::V5::*sdeB* Sh cassette. In this construct, the *sdeB*$^{Af}$ was placed under the control of the *gpdA* promoter and fused to an N-terminal V5 epitope tag, while retaining the *sdeB* 3' regulatory region. Primers used for amplification and assembly are indicated in the scheme. Both cassettes shown in A and B are flanked by the 5' and 3' Sh homology arms to allow targeted integration at the *aft4* Sh locus. (C) Strategy used for the generation of the Δ*sdeB* deletion cassette. Independent fragments corresponding to the 5' and 3' flanking regions (~1.5 kb) of the *sdeB* locus and the *ptrA* selectable marker (2.025 kb) were amplified by PCR using the primers indicated in the figure and assembled in *S. cerevisiae*. The top scheme shows the wild-type *sdeB* locus, and the bottom scheme shows the expected recombinant Δ*sdeB* locus after replacement. (D) PCR-based validation of the Δ*sdeB* and *xylP::sdeA* Δ*sdeB* mutants after transformation. (E) Southern blot validation of the mutants. Genomic DNA was digested with *Eco*RI and hybridized using the 5' flanking region as probe (green line). The expected hybridizing fragments for the wild-type and mutant loci are indicated in blue and red, respectively. (F) PCR validation of the complemented strain Δ*sdeB::sdeB*$^{+}$ after transformation of the Δ*sdeB* mutant. Primers IM-639 and IM-517 were used to amplify the indicated 2.097-kb fragment, confirming successful complementation.
